# Catabolism of extracellular glutathione is a driver of metastasis formation

**DOI:** 10.64898/2026.10.12.757337

**Authors:** Felix C. E. Vogel, Florian Röhrig, Danny Schilling, Adriano B. Chaves-Filho, Greta Mattavelli, Maximilian Krämer, Julie Haenlin, Alina M. Winkelkotte, Valentine Fournier, Kamal M. Al-Shami, Hannah Rohdjeß, Philipp Pöller, Tanja Poth, Christoph Stange, Bernhard Küster, Lukas Wagener, Verena Thewes, Andreas Schneeweiss, Peter Lichter, Lisa Schlicker, Marteinn T. Snaebjornsson, Angela Riedel, Almut Schulze

## Abstract

Metastasizing cancer cells require metabolic plasticity to adapt to the microenvironment of the target organ. Analyzing metabolites in interstitial fluids, we found that the liver microenvironment is devoid of cystine but contains high concentrations of glutathione (GSH). Detailed mechanistic analysis revealed that cancer cells degrade extracellular GSH utilizing γ-glutamyl-transferase (GGT) activity for provision of cysteine to support proliferation and prevent ferroptosis under cystine-free conditions. This was independent of system Xc^-^ but required activity of glutamate-cysteine ligase (GCLC). GGT1 expression was associated with poor survival and upregulated in liver metastases of human breast cancer patients. Overexpression of Ggt1 in murine breast cancer cells enhanced their capacity for liver colonization and promoted lung metastases after orthotopic implantation. This identifies extracellular GSH catabolism as driver of metastasis that supports proliferation and oxidative stress resistance in metabolically challenging microenvironments.

## Introduction

Metastasis remains the primary cause of cancer-related deaths (*1*). Disseminated cancer cells require metabolic flexibility and plasticity to adapt to the changing microenvironmental conditions encountered along the metastatic cascade (*2*). Targeting metabolic processes that enable cancer cells to survive during metastatic dissemination may thus provide novel therapeutic options. Metastatic colonization of specific organs, called tropism, is a complex process that is determined by multiple factors (*3–5*) and organ-specific metabolic adaptations have been identified in multiple contexts (*6–10*). In addition, cancer cells are exposed to elevated oxidative stress during metastasis formation (*11–14*). Oxidative stress causes lipid damage that leads to ferroptosis, an iron-dependent form of non-apoptotic cell death (*15*). Glutathione peroxidase 4 (GPX4) prevents ferroptosis by converting lipid peroxides into non-toxic alcohols (*15–19*). This reaction requires reduced glutathione (GSH), making GSH a rate-limiting factor for ferroptosis prevention. Synthesis of GSH depends on the availability of cysteine, a conditionally essential amino acid required for protein synthesis and the generation of coenzyme A and iron-sulfur clusters (*20, 21*). Most cancer cells satisfy their cysteine requirement through cystine uptake in exchange for glutamate via system Xc-(SLC7A11/ SLC3A2) (*22–24*). Cystine deprivation should thus impair cancer cell viability and proliferation in the absence of alternative sources. Catabolism of extracellular GSH by γ-glutamyl-transpeptidases (GGT) (*25, 26*) has been shown to confer resistance to experimental cystine deprivation in glioma cells *in vitro*(*27*) and to accelerate growth and cisplatin resistance of prostate cancer cells *in vivo* (*28*). Recently, it was shown that experimental overexpression of GGT1 in cancer cells restored proliferation and oxidative stress resistance in cystine-depleted environments *in vitro* and promoted xenograft tumor formation *in vivo* (*29*).

To identify metabolic determinants of metastasis formation, we conducted quantitative analysis of metabolites in interstitial fluids from different metastatic target organs and established that the liver is devoid of cystine but rich in extracellular GSH. Catabolism of extracellular GSH emerged as an essential process that provides cancer cells with cysteine to support proliferation and oxidative stress resistance in cystine-deprived conditions. Furthermore, overexpression of Ggt1 in breast cancer cells enhanced metastatic liver colonization and promoted spontaneous metastasis in an orthotopic model. Our findings identify catabolism of extracellular GSH as an alternative source of cysteine and suggest that targeting GGT restricts the ability of cancer cells to adapt to cystine-low microenvironments.

## Results

### GSH in liver interstitial fluid supports cancer cell proliferation and survival

In order to analyze the metabolic microenvironment experienced by metastatic cancer cells during the colonization of different organs, we extracted plasma and interstitial fluids (IF) from lungs and livers of healthy mice (*30, 31*) and subjected them to liquid chromatography coupled mass spectrometry (LC-MS) (**Fig. 1A**). To account for matrix effects, we used a standard addition method (SAM) by spiking fixed amounts of fetal bovine serum (FBS) with increasing amounts of Mouse Plasma-like Medium (MPLM) formulated based on metabolite concentrations of mouse plasma (*32, 33*). Absolute metabolite concentrations for liver-IF, lung-IF and plasma were then calculated from equal volumes of FBS analyzed in parallel (**fig. S1A and B**).

**Figure 1:**
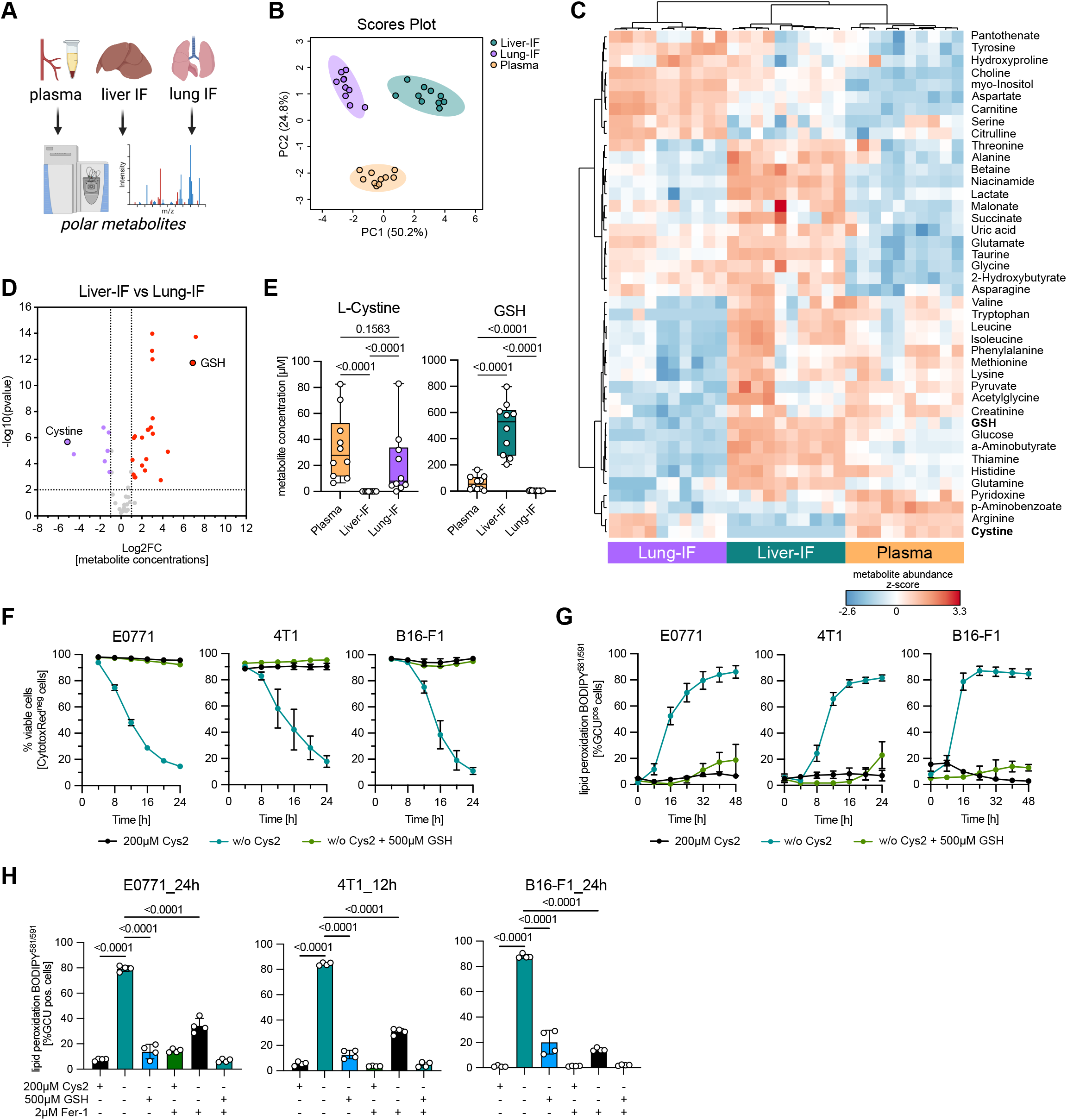
Quantification of organ-specific metabolic microenvironments identifies GSH as critical factor for cell viability and inhibition of oxidative stress. **(A)** Diagram of extraction strategy to obtain liver and lung interstitial fluid (IF) together with plasma from healthy BALB/c mice for LC-MS analysis. Created in https://BioRender.com. **(B)** Principal component analysis (PCA) of polar metabolite abundances in the different fluids (n=10). **(C)** Heatmap of 42 significantly altered metabolites derived from absolute quantifications in the different biological fluids (n=10). GSH and cystine are highlighted. **(D)** Volcano plot comparing metabolite abundance between liver-IF and lung-IF. **(E)** Absolute metabolite concentrations of cystine and GSH in the different fluids. Data are displayed as box and whisker plots, showing mean, 25^th^ and 75^th^ percentile and min-max. Significance was calculated using one-way ANOVA with Tukey’s post-hoc test on log transformed data (n=10). (F) Cell viability of E0771, 4T1 and B16-F1 cells cultured under the indicated conditions determined by CytotoxRed staining and live cell imaging (Incucyte) over 24 hours. Data show mean ± SD of biologically independent samples (n=4). **(G)** Induction of lipid peroxidation in E0771, 4T1 and B16-F1 cells cultured under the indicated conditions determined by BODIPY_581/591_-C_11_ staining and live cell imaging followed by AI cell-by-cell analysis (Incucyte) over 24 (4T1) or 48 (E0771 and B16F1) hours. Data show mean ± SD of biologically independent samples (n=4). **(H)** 4T1, E0771, 4T1 and B16-F1 cells were cultured with or without cystine (Cys2) or with addition of 500 µM GSH either in the presence or absence of 2 µM ferrostatin (Fer-1). Lipid peroxidation was determined by BODIPY_581/591_-C_11_ staining and live cell imaging followed by AI cell-by-cell analysis (Incucyte) over 12 (4T1) or 24 (E0771 and B16F1) hours. Data show mean ± SD of biologically independent samples (n=4).

Principal component analysis (PCA) showed clear discrimination of the different fluids (**Fig. 1B**), with 42 of 47 metabolites showing significant differences (**Fig. 1C and Data S1**). Direct comparison of metabolite levels between liver-IF and lung-IF revealed a marked reduction in cystine in liver-IF, while reduced glutathione (GSH) was strongly increased in liver-IF (**Fig. 1D and E**). Concentrations for cystine in lung-IF and plasma ranged up to 83 µM, while it remained below the limit of detection in all samples of liver-IF (**Fig. 1E**). Conversely, while GSH was mostly undetectable in lung-IF, concentration of this metabolite in liver-IF ranged from 203 to 797 µM (**Fig. 1E**). As expected from the physiological function of the liver in metabolic homeostasis, levels of glucose and lactate, as well as the amino acids glutamine, alanine, leucine, and taurine were strongly enriched in liver-IF (**fig. S1C and D**).

Despite being a non-essential amino acid, cysteine is considered to be rate-limiting for proliferation, as it is required for multiple cellular functions, including protein synthesis and redox control (*20, 34*). Cysteine is generally taken up in its oxidized form as cystine via system Xc-, which imports cystine in exchange for glutamate (*23, 24*). Based on our data, we considered that metastasizing cancer cells would be exposed to a cystine-low environment during liver colonization. Culture of 4T1 murine breast cancer cells in different concentrations of cystine or cysteine caused a severe reduction in cell number at concentrations below 30 µM for cystine and 60 µM for cysteine (**fig. S2A**). Addition of the lipophilic radical trapping antioxidants ferrostatin-1 (Fer-1) or liproxstatin-1 (Lip-1), the iron-chelator deferoxamin (DFO), or the antioxidant N-Acetyl-L-Cysteine (NAC) restored 4T1 and E0771 cell viability in the absence of cystine. In contrast, Necrostatin-2 (Nec-2) or Z-VAD-FMK did not restore viability in this condition, excluding major contribution by necroptosis or apoptosis (**fig. S2B to E**).

We next supplemented cystine-free medium with GSH at concentrations comparable to those detected in liver-IF (250 or 500 µM) and found that this restored cell number and proliferation in murine breast cancer (4T1, E0771) and melanoma cells (B16-F1) (**fig. S3A**). Time resolved analysis showed a loss of viability and proliferation in cystine-free medium over 24 hours, which was prevented by addition of 500 µM GSH (**Fig. 1F and fig. S3B**). A similar effect on cell viability was also observed in human breast cancer (MDA-MB-231, MDA-MB-436 and MDA-MB-468) and melanoma cells (A375) (**fig. S3C**). Morphological analysis of cells cultured in cystine-free medium revealed cytoplasmic swelling and leakage typically associated with ferroptosis and these phenotypes were efficiently prevented by addition of GSH (**fig. S3D**). Furthermore, GSH also efficiently abolished the induction of lipid peroxidation, a hallmark of ferroptosis (*18*), in cystine-free medium (**Fig. 1G and fig. S3E**). The inhibitory effect of GSH on lipid peroxidation in cystine-depleted conditions was similar or even stronger than that of ferrostatin (**Fig. 1H**), confirming that extracellular GSH can substitute for cystine in preventing oxidative stress to support cancer cell proliferation and survival.

### Cancer cells degrade extracellular GSH to obtain substrates for intracellular GSH synthesis

Extracellular GSH can be cleaved by γ-glutamyl-transpeptidases (GGT) to produce cysteinyl-glycine and either free glutamate or a new γ-glutamyl-compound if a suitable acceptor, i.e. another amino acid or dipeptide, is present (*26*). The resulting cysteinyl-glycine can either be imported or cleaved into cysteine and glycine, which are taken up via dedicated transporters (**Fig. 2A**).

**Figure 2:**
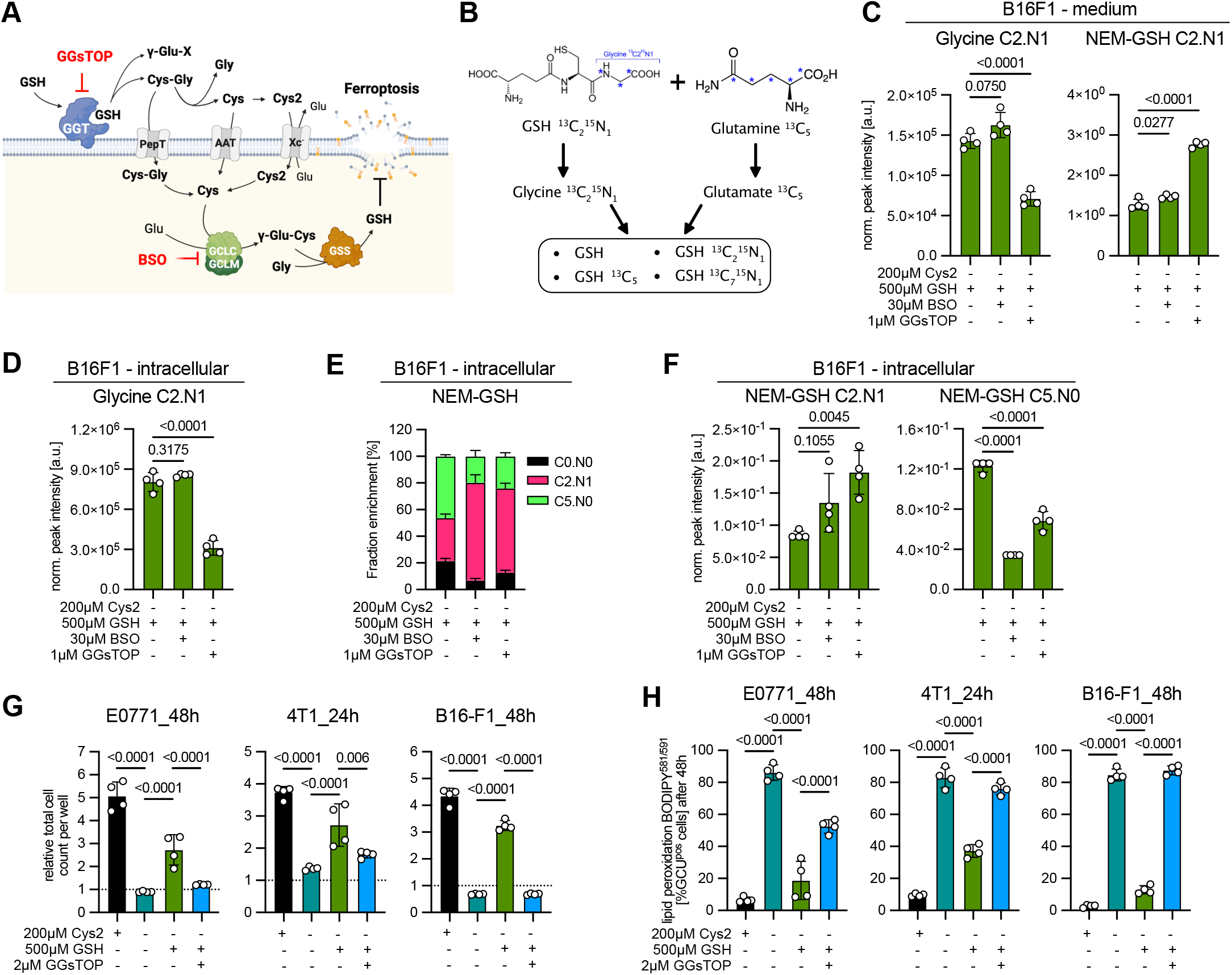
Cancer cells degrade extracellular GSH to generate substrates for *de novo* GSH synthesis. (**A**) Diagram indicating processes involved in the degradation of extracellular GSH and intracellular *de novo* GSH synthesis. Inhibitors of essential components of the pathway are also indicated. Created in https://BioRender.com. **B)** Schematic overview of the double labelling strategy showing the resulting isotopologues of GSH. (**C**) B16F1 cells were pre-labelled in glutamine-free RPMI medium supplemented with 2 mM L-Glutamine-^13^C_5_ for 24 hours, then switched to cystine/glutamine-free medium supplemented with 500 µM GSH-^13^C_2_^15^N_1_ and 2 mM L-glutamine-^13^C_5_ and treated with 30 µM BSO or 1 µM GGsTOP. Samples were taken after 24 hours and relative abundance of Glycine-C2.N1 and NEM-GSH C2.N1 in medium was quantified after NEM-derivatization. Data show mean ± SD of biologically independent samples (n=4). Significance was calculated using one-way ANOVA with Dunnett’s post-hoc test. (**D**) Abundance of intracellular Glycine-1^3^C_2_.^15^N_1_ in cells from C. (**E**) Fractional labelling of intracellular GSH. (**F**) Abundance of intracellular GSH-^13^C_2_^15^N_1_ and GSH-^13^C_5_^15^N_0_ In the same cells. (**G**) E0771, 4T1 and B16-F1 cells were cultured in RPMI containing 200 µM cystine (Cys2) or in cystine-free conditions supplemented with or without 500 µM GSH for 24 (4T1) or 48 (E0771 and B16F1) hours. GGT activity was blocked by addition of 2 µM GGsTOP. Cell number was determined by automated image analysis (Incucyte). Data show mean ± SD of biologically independent samples (n=4). Significance was calculated using one-way ANOVA with Tukey’s post-hoc test. (**H**) Lipid peroxidation in cells treated as in G was determined by BODIPY^581/591^ C11 staining at indicated time points. Data show mean ± SD of biologically independent samples (n=4). Significance was calculated using one-way ANOVA with Tukey’s post-hoc test.

To obtain mechanistic detail of extracellular GSH catabolism, we analyzed culture medium and extracts of cancer cells exposed to different culture conditions using LC-MS. Time-resolved analysis of supernatants of B16-F1 cells showed accumulation of γ-glutamyl-alanylglycine, a tripeptide produced by the transpeptidase activity of GGT only in presence of GSH (**fig. S4A**). In 4T1 cells, addition of extracellular GSH to cystine-free medium restored half of the intracellular GSH pool compared to cystine-containing conditions (**fig. S4B**). This was accompanied by an increase in cysteinyl-glycine both in medium and in cells (**fig. S3C**).

Using stable isotope tracing, we observed that addition of GSH carrying labelled carbon and nitrogen atoms in the glycine moiety (GSH-^13^C_2_^15^N_1_) to E0771 cells cultured in cystine-free medium resulted in a time-dependent release of labelled glycine, indicative of GGT activity (**fig. S4D)**. Analysis of intracellular metabolites revealed fractional labelling of the GSH pool to approx. 15% after 24 hours, while glycine was labelled to a much lesser degree (**fig. S4E**). This suggests that a substantial proportion of intracellular GSH must be derived either from uptake of intact GSH or from uptake of cysteinyl-glycine, which then might serve as a substrate for GSH synthesis.

To distinguish between these two options, we applied stable isotope tracing strategies. First, we cultured E0771 and 4T1 cells in the presence of GSH and L-glutamine-^13^C_5_ to determine contribution of extracellular GSH to intracellular GSH synthesis. Fractional labelling showed that approx. 30% of intracellular GSH is synthesized from ^13^C_5_-glutamine (**fig. S4F**). Treatment with the γ-glutamyl-cysteine ligase (GCLC) inhibitor buthionine sulphoximine (BSO) or addition of GGsTOP, a highly potent irreversible GGT inhibitor (*35*), resulted in a strong reduction of the GSH M+5 pool and fractional labelling, suggesting that GGT activity is required to generate substrates for *de novo* GSH synthesis (**fig. S4F and G**). GGsTOP also reduced the depletion of GSH from the culture medium and blocked the accumulation of γ-glutamyl-alanylglycine observed after 12 hours in B16-F1 cells (**fig. S4H**).

We next applied a double labelling strategy to reveal the impact of GGT inhibition on *de novo* GSH synthesis (**Fig. 2B**). B16-F1 cancer cells were first cultured in medium containing L-glutamine-^13^C_5_ for 24 hours before medium was changed to cystine-free medium containing ^13^C_5_-glutamine together with 500 µM of GSH-^13^C_2_^15^N_1_. Cells were incubated for an additional 24 hours and metabolites arising from these labelled precursors were analyzed. Accurate quantification of thiol-containing metabolites was achieved by applying N-ethylmaleimide (NEM) derivatization (*36, 37*). In medium, treatment with GGsTOP, but not BSO, reduced the abundance and fractional enrichment of C2.N1-labelled glycine while the abundance of C2.N1-labelled GSH was increased, confirming GSH catabolism by GGT (**Fig. 2C and fig. S5A)**. Analysis of intracellular metabolites showed that GGsTOP efficiently reduced levels and fractional enrichment of C2.N1-labelled glycine (**Fig. 2D and fig. S5B**), demonstrating that GSH catabolism contributes to the intracellular glycine pool. Analysis of fractional enrichment intracellular GSH (**Fig. 2E**) showed that approximately 50% is represented by the C5.N0 isotopologue derived from de novo synthesis using labelled glutamine, while approximately 30% was present as C2.N1 isotopologue, most likely derived from uptake of the intact molecule. The C7.N1 isotopologue derived from de novo synthesis using labelled glutamine and glycine released from GSH catabolism was not detected, as C2.N1 glycine only contributed 5% to the total glycine pool (**fig. S5B**). Treatment with GGsTOP or BSO reduced abundance and fractional labelling of C5.N0-GSH (**Fig. 2F and fig. S5C)**. Conversely, abundance and enrichment of C2.N1-labelled GSH was increased, suggesting that GSH uptake compensates for inhibition of GSH catabolism and synthesis (**Fig. 2E and F, fig. S5C**). As a consequence, intracellular GSH levels were reduced by BSO but not after GGT inhibition (**fig. S5D)**. No changes were observed in the labelling of intracellular L-Glutamine and L-Glutamate, which were both labelled to a high degree (**fig. S5E**).

We also established the effect of GGT inhibition on cell viability and oxidative stress. While the inhibitor had no adverse effect on cells cultured in cystine-containing medium (**fig. S5F and G**), it efficiently blocked GSH-dependent proliferation and inhibition of lipid peroxidation in cystine-free medium (**Fig. 2G and H**). Similar results were obtained in human breast cancer and melanoma cells, where ferroptotic cell death was confirmed by Lip-1 rescue (**fig. S5G**). These results indicate that in addition to uptake of the intact molecule, catabolism of extracellular GSH by GGT provides substrates for intracellular GSH synthesis to support cell viability and prevent oxidative stress in cancer cells.

### Extracellular GSH catabolism provides cells with cysteine independent of Xc^-^

To determine the mechanism by which products of extracellular GSH catabolism are taken up by cancer cells (**Fig. 3A**), we treated cells with erastin, a selective inhibitor of Xc**⁻** (*38*). Erastin caused a strong reduction of cell number in cystine-containing medium but only had a minimal effect in cystine-depleted condition supplemented with GSH (**Fig. 3B and fig. S6A**), suggesting that system Xc**⁻** is dispensable under this condition. Next, we substituted GSH with an equimolar concentration of cysteinyl-glycine (Cys-Gly), the dipeptide released by GGT-dependent cleavage of GSH. This showed that Cys-Gly, but not glycyl-glycine (Gly-Gly), fully restored proliferation in the absence of cystine (**Fig. 3C and fig. S6B**). Cys-Gly also rendered cells fully insensitive to GGT inhibition (**fig. S6C**), while inhibition of GCLC with BSO still abolished cell viability (**fig. S6D**). However, cell viability rescue by Cys-Gly was highly dependent on Xc^-^, as it was fully blocked by erastin (**Fig. 3D**).

**Figure 3:**
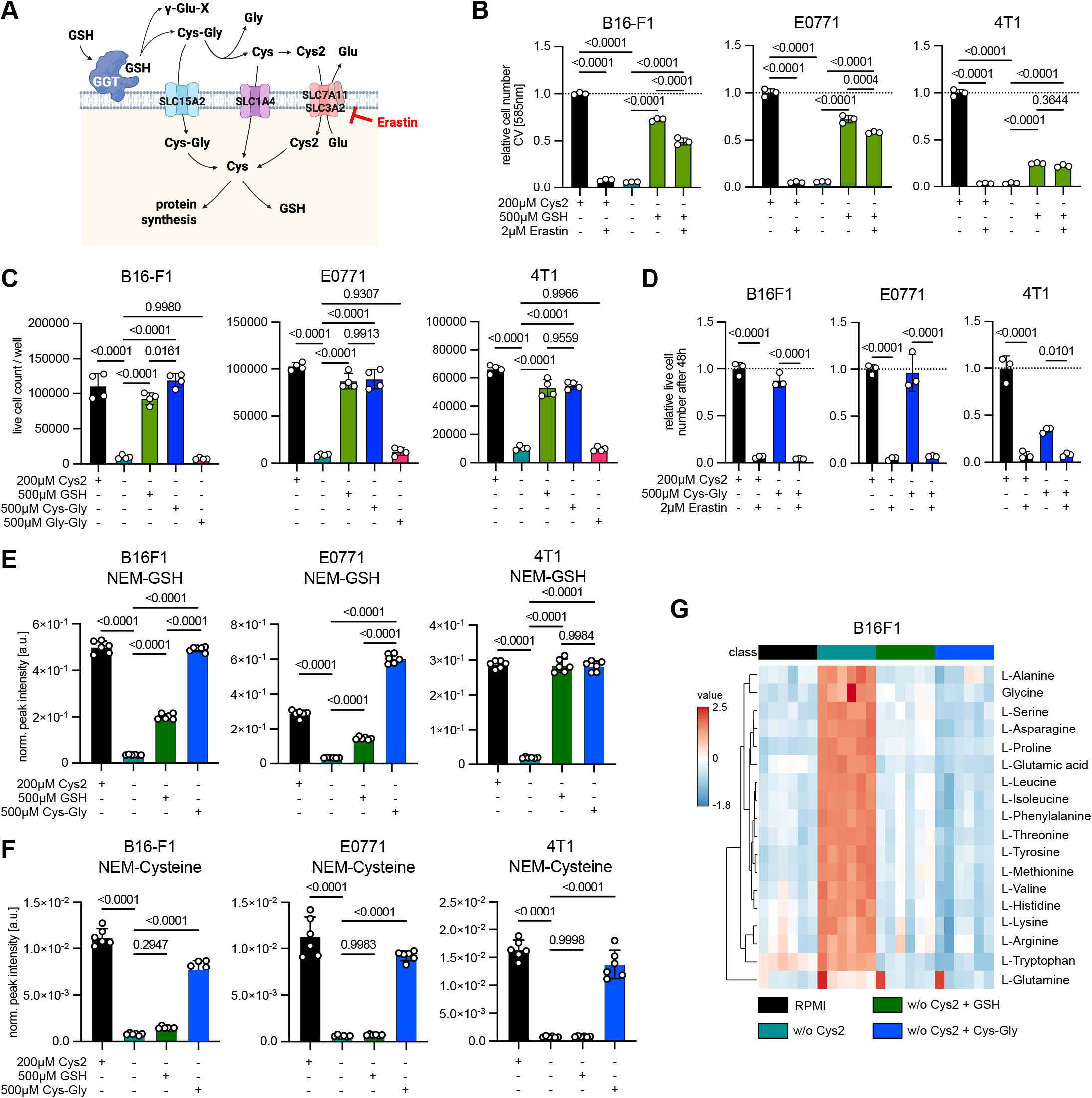
Extracellular GSH protects cancer cells against ferroptosis independent of system Xc^-^. (**A**) Diagram showing different transporters mediating uptake of products of GSH catabolism. Created in https://BioRender.com. (**B**) B16-F1, E0771 and 4T1 cells were cultured in RPMI (200 µM Cys2) or in cystine-free medium in the presence or absence of 500 µM GSH with or without addition of 2 µM erastin for 48 hours. Relative cell number was determined by crystal violet staining. Data show mean ± SD of biologically independent samples (n=3). Significance was calculated using one-way ANOVA with Tukey’s post-hoc test. (**C**) The same cells were cultured in RPMI (200 µM Cys2) or in cystine-free medium with or without addition of 500µM GSH or equimolar concentration of cysteinyl-glycine (Cys-Gly) or glycinyl-glycine (Gly-Gly) for 48 hours. Number of live cells per well was determined by automated imaging (Incucyte). Data show mean ± SD of biologically independent samples (n=4). Significance was calculated using one-way ANOVA with Tukey’s post-hoc test. (**D**) Cells were cultured in RPMI (200 µM Cys2) or cystine-free medium in the presence of 500 µM cysteinyl-glycine (Cys-Gly) and treated with 2 µM Erastin for 48h. Number of live cells per well was determined by automated imaging (Incucyte). Data show mean ± SD of biologically independent samples (n=3). Significance was calculated using one-way ANOVA with Tukey’s post-hoc test. (**E**) Cells were cultured in RPMI (200 µM Cys2) or in cystine-free medium in the presence or absence of 500 µM GSH or 500 µM cysteinyl-glycine (Cys-Gly) for 6 hours. Thiol-metabolites were detected by derivatization with N-ethylmaleimide (NEM) followed by LC-MS. Graph shows intracellular levels of NEM-GSH. Data show mean ± SD of biologically independent samples (n=6). Significance was calculated using one-way ANOVA with Tukey’s post-hoc test. (**F**) Intracellular levels of NEM-Cysteine in the same cells as in E. (**G**) Heatmap showing relative abundance of amino acids in B16-F1 cells treated as in E.

We conclude that the high abundance of extracellular GSH might retain free cysteine in its reduced form, to make it available for uptake by other amino acid transporters, such as SLC1A4 (ASCT1). This is supported by data from the Cancer Therapeutics Response Portal (*39*) correlating expression of the cysteine transporter *SLC1A4* with resistance to GPX4 inhibitors across a large number of cell lines (**fig. S4E**). Nevertheless, it is also possible that direct uptake of Cys-Gly contributes to cell viability, as expression of *SLC15A2*, the dipeptide transporter mediating its uptake (*40*), also correlated with ferroptosis resistance (**fig. S4E**).

To obtain further insight, we analyzed intracellular metabolites of cells exposed to the different conditions for 6 hours. This revealed that addition of Cys-Gly was more efficient than GSH in restoring intracellular GSH and GSSG levels in cystine-depleted conditions in B16-F1 and E0771 cells (**Fig. 3E and fig. S7A**). Moreover, only Cys-Gly increased intracellular cysteine levels (**Fig. 3F**), while intracellular levels of cystine remained low under all cystine-deplete conditions (**fig. S7B**). GSH was also less efficient than Cys-Gly in preventing the accumulation of γ-glutamyl-glycine, a dipeptide produced by a non-canonical activity of GCLC during cystine starvation (*41*) (**fig. S7C**). Furthermore, cells exposed to GSH under cystine starvation showed elevated sensitivity towards BSO (**fig. S7D**), suggesting that GCLC inhibition is highly toxic under these conditions. While this was initially puzzling, we reasoned that intracellular cysteine might be rapidly metabolized as it is required for multiple processes, including protein biosynthesis (*20*). Indeed, cystine depletion resulted in a strong accumulation of all other amino acids, suggesting that protein synthesis is inhibited due to the lack of cysteine. Interestingly, addition of GSH or Cys-Gly relieved amino acid accumulation in cystine-free conditions, indicating that protein synthesis was restored (**Fig. 3G and fig. S7E and F**).

This suggests that the catabolism of extracellular GSH is an inefficient process that provides relatively small amounts of free cysteine, which are rapidly taken up and used by cancer cells. Nevertheless, extracellular GSH catabolism is sufficient to produce sufficient cysteine to restore proliferation in cystine-depleted conditions.

### Ggt1 contributes to GSH catabolism to support cell survival in cystine-free environments

Mammalian genomes encode at least four γ-glutamyl-transpeptidase enzymes (*GGT1*, *GGT5*, *GGT6* and *GGT7*) (*25*). Only GGT1 and GGT5 have GSH hydrolase activity (*42*), while GGT7 may protect from oxidative stress by an unknown mechanism (*43*). *GGT1*, *GGT5* and *GGT7* were expressed in human breast cancer cell lines, but only expression of GGT1 correlated with GGT activity (**fig. S8A and B**). Ggt1 expression also correlated with GGT activity in murine cancer cells, with highest levels found in B16-F1 (**Fig. 4A and B and fig. S8C**).

**Figure 4:**
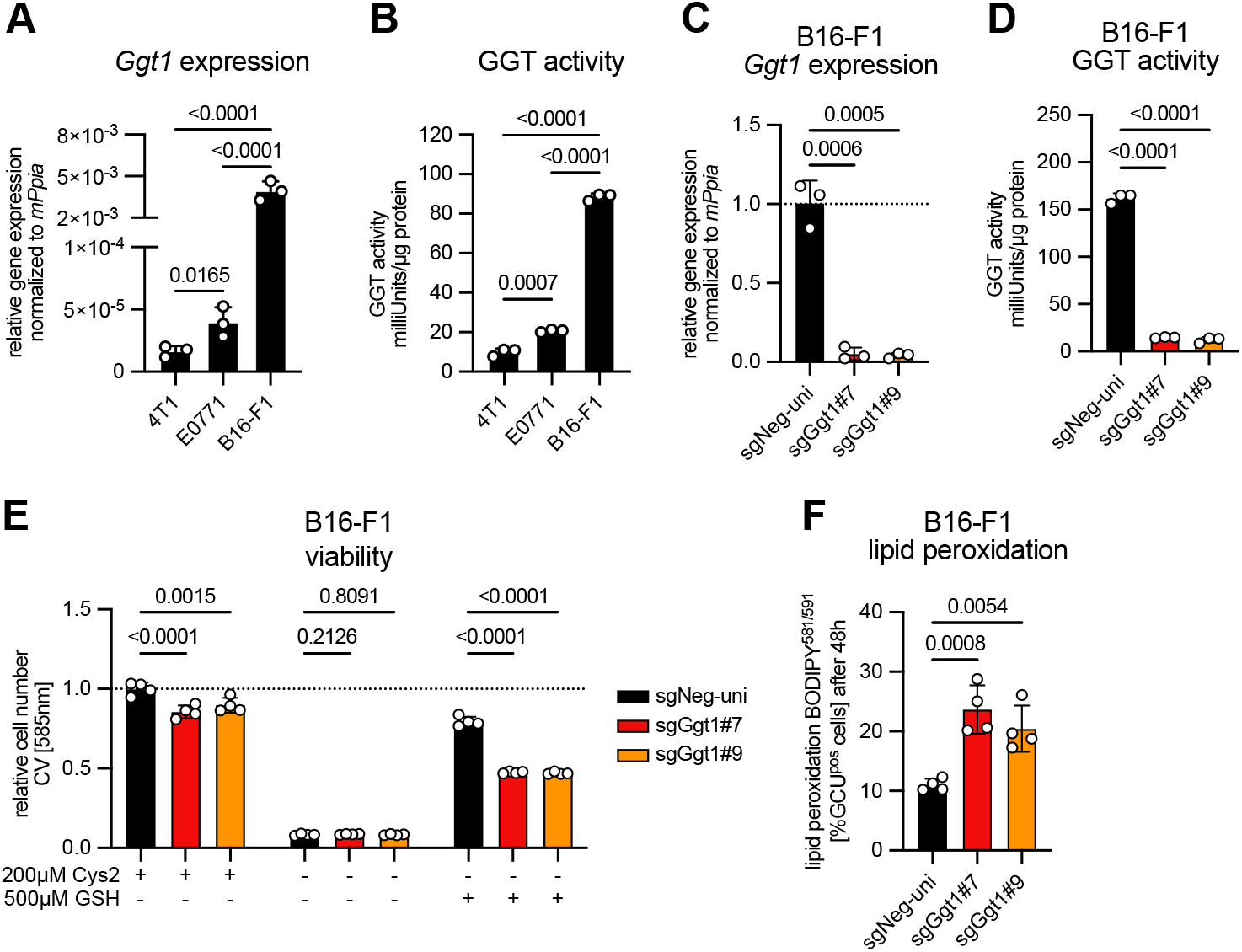
Depletion of Ggt1 attenuates extracellular GSH catabolism and viability rescue in cystine-free environments. **A)** Relative expression of *Ggt1* mRNA in 4T1, E0771 and B16-F1 cells normalized to peptidylprolyl isomerase A (*Ppia*). Data show mean ± SD of biologically independent samples (n=3). Significance was calculated using one-way ANOVA with Tukey’s post-hoc test on log transformed data. (**B**) Enzymatic GGT activity in extracts from the same cells. Data show mean ± SD of biologically independent samples (n=3). Significance was calculated using one-way ANOVA with Tukey’s post-hoc test. (**C**) Relative expression of *Ggt1* mRNA in B16-F1 control and Ggt1 knockout cells. Data show mean ± SD of biologically independent samples (n=3). Significance was calculated using one-way ANOVA with Dunnett’s post-hoc test on log transformed data. (**D**) GGT activity in the same cells. Data show mean ± SD of biologically independent samples (n=3). Significance was calculated using one-way ANOVA with Dunnett’s post-hoc test. (**E**) B16-F1 control and Ggt1 knockout cells were cultured in RPMI or cystine-free medium in the presence or absence of 500 µM GSH for 48 hours. Relative cell numbers were determined by crystal violet staining. Data show mean ± SD of biologically independent samples (n=4). Significance was calculated using two-way ANOVA with Dunnett’s post-hoc test on log transformed data. (**F**) B16-F1 control and Ggt1 knockout cells were cultured in cystine-free medium in the presence of 500 µM GSH for 48 hours. Lipid peroxidation was determined by BODIPY^581/591^-C11 staining. Data show mean ± SD of biologically independent samples (n=4). Significance was calculated using one-way ANOVA with Dunnett’s post-hoc test.

Deletion of Ggt1 in B16-F1 cells abolished expression and enzymatic activity but had no effect on proliferation in cystine-containing medium (**Fig. 4C and D and fig. S8D**). Ggt deletion also lowered the capacity of cells to catabolize extracellular GSH, demonstrated by stable isotope tracing (**fig. S8E and F**). Furthermore, the ability of extracellular GSH to prevent amino acid accumulation in cystine-free conditions was reduced in Ggt1 knockout cells (**fig. S8G**). Consequently, Ggt1 deletion reduced the ability of extracellular GSH to restore proliferation and prevent lipid peroxidation in cystine-free conditions (**Fig. 4E and F and fig. S8H**). This confirms that Ggt1 contributes to extracellular GSH catabolism to prevent oxidative stress and support proliferation. However, in contrast to the effects of chemical inhibition by GGsTOP, the effects were less pronounced, suggesting contribution by other isoforms or alternative mechanisms.

### Overexpression of Ggt1 enhances GSH catabolism and drives liver metastasis

It has been shown that murine breast cancer cells metastasizing to the liver increase GSH synthesis (*44*) and that depletion of GCLC reduces the ability of human breast cancer cells to form metastases in this organ (*44*). We therefore considered that GGT-dependent catabolism of extracellular GSH could prevent oxidative stress during liver colonization and outgrowth.

Analysis of TCGA data revealed that high expression of *GGT1* is associated with poor survival in breast, pancreatic and lung squamous cancers (**fig. S9A**). Furthermore, *GGT1* expression was increased in patients with stage III or IV breast cancers and showed a strong link with poor survival in this patient cohort (**fig. S9B and C**). Analysis of data from 16 primary breast cancers and 67 matched metastases from different sites (*45*) showed that expression of *GGT1*, but not other GGT enzymes, is significantly increased in liver metastases compared to primary tumors (**Fig. 5A and fig. S9D**). This was even more pronounced when comparing primary tumors and liver metastases from the same patient (**fig. S9E**). Liver metastases also showed upregulation of genes mapping to GSH metabolism and oxidative stress (**fig. S9F**), suggesting a higher demand for GSH in metastases.

**Figure 5:**
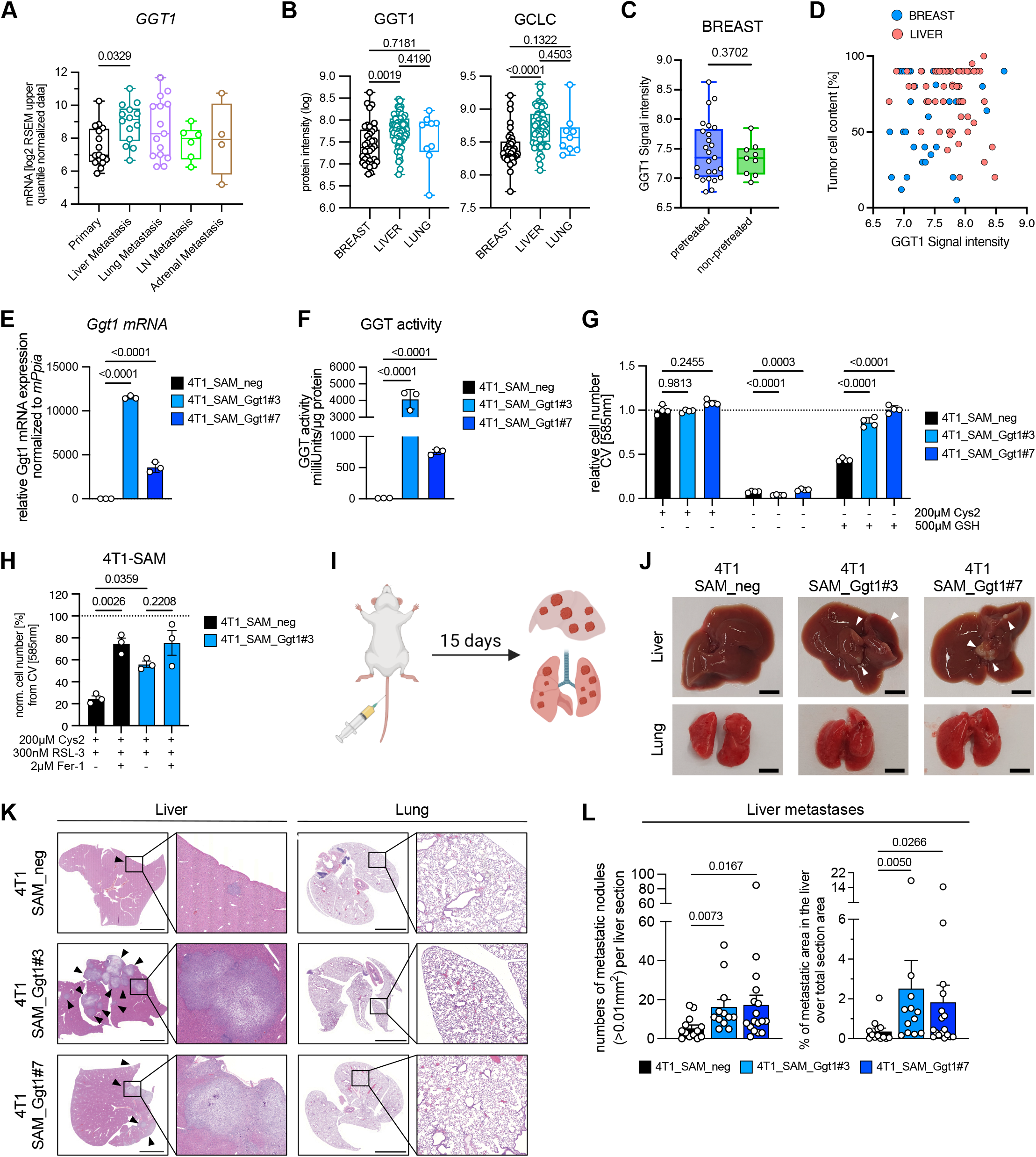
Catabolism of extracellular GSH drives colonization and outgrowth in the liver microenvironment. (**A**) Expression of *GGT1* mRNA in a panel of TNBC primary tumors and metastases from multiple sites from GSE110590 (*45*). Box and whisker plots display median, 25^th^ and 75^th^ percentile and min-max. Significance was calculated using one-way ANOVA with Dunnett’s post-hoc test (primary n=16, liver metastasis n=14; lung metastasis n=15; lymph node (LN) metastasis n=6; adrenal metastasis n=4). (**B**) GGT1 and GCLC protein intensity in primary tumors as well as liver and lung metastases from breast cancer patient samples collected within the CATCH and COGNITION registry trials (NCT05652569 and NCT05906407) analyzed by quantitative proteomics. Box and whisker plots display median, 25^th^ and 75^th^ percentile and min-max. Statistical significance was calculated using one-way ANOVA with Tukey’s post-hoc test (breast n=36 for GGT1 and breast n=37 for GCLC; liver n=60; lung n=9). (**C**) Comparison of breast tumors from patients with and without prior treatment. Statistical significance was calculated using Welch’s t test (pretreated n=25; non-retreated n=9). Treatment modalities are listed in Extended Data Table 2. (**D**) Correlation of GGT1 signal intensity and tumor cell content for breast and liver samples from the same study (R^2^: breast=0.027; liver=0.043). (**E**) Expression of *Ggt1* mRNA after CRISPR/SAM-mediated overexpression in 4T1 cells using two independent guide RNAs (sgGgt1#3 and sgGgt1#7) compared to non-targeting control (sgNeg1) normalized to *Ppia*. Data show mean ± SD of biologically independent samples (n=3). Significance was calculated using one-way ANOVA with Dunnett’s post-hoc test on log transformed data. (**F**) GGT activity in 4T1 cells after CRISPR/SAM-mediated overexpression of Ggt1 determined by enzymatic assay. Data show mean ± SD of biologically independent samples (n=3). Significance was calculated using one-way ANOVA with Dunnett’s post-hoc test. (**G**) 4T1 control and Ggt1 overexpressing cells were cultured in complete or cystine-free medium in the presence or absence of 500 µM GSH for 48 hours. Relative cell numbers were determined by crystal violet staining. Data show mean ± SD of biologically independent samples (n=4). Significance was calculated using two-way ANOVA with Dunnett post-hoc test on log transformed data. (**H**) Control and Ggt1 overexpressing cells were cultured under the indicated conditions and treated with 300 nM of the ferroptosis inducer RSL3 either in the presence or absence of 2 µM ferrostatin (Fer-1). Viability was determined by automated cell imaging (Incucyte). Data show mean ± SD of biologically independent samples (n=4). (**I**) Diagram indicating multi-organ metastasis formation after hydrodynamic tail vein (HDTV) injection of 4T1 cells. Created in https://BioRender.com. (**J**) Representative images of livers and lungs from mice injected with 4T1 control (SAM_neg) and Ggt1 overexpressing cells (SAM_Ggt1#3 and SAM_Ggt1#7). Arrowheads indicate visible metastases in liver. Scale bars = 5mm. (**K**) Representative histological sections of matched liver and lung tissues of mice after HDTV injection of control or Ggt1-overexpressing 4T1 cells (3 x 10^5^ cells per injection). Arrowheads indicate metastatic nodules visible in liver sections; no metastatic nodules were detected in the lungs of mice in all groups. Scale bar 400 µm. (**L**) Quantification of the number of metastatic nodules and metastatic area in livers from mice after HDTV injection of control or Ggt1-overexpressing 4T1 cells. Data are shown as mean ± SD of each cohort (sgNeg1 n=13; sgGgt1 n=12; sgGgt1#7 n=17). Significance was calculated using one-way ANOVA with Dunnett’s post-hoc test on log transformed data.

We next investigated samples from two ongoing registry trials aimed at genomic profiling of neoadjuvant chemotherapy-resistant tumors in high-risk breast cancer patients (NCT05652569 and NCT0590640). As these cohorts represent advanced disease, many patients have been pre-treated with chemotherapy and targeted agents (**Table S1**). Quantitative proteomics revealed increased expression of GGT1 and GCLC in extracts from liver metastases compared to breast tissues (**Fig. 5B**), suggesting that GSH catabolism is relevant in this setting. Differential expression in lung metastases did not reach statistical significance due to small sample size. Importantly, we observed no difference in GGT1 expression in breast tumors with or without pre-treatment, confirming that the increased expression in liver metastasis was not caused by therapy-induced oxidative stress (**Fig. 5C**). Tumor cell content showed no association with GGT1 expression in breast and liver samples (**Fig. 5D**), excluding sampling effects.

This suggests that extracellular GSH catabolism by GGT1 could promote dissemination and outgrowth of breast cancer cells in the liver microenvironment. We therefore transduced 4T1 cells with gRNAs targeting CRISPR/sgRNA-directed synergistic activation mediator (SAM) system (*46*) to the *Ggt1* promoter. This resulted in strong upregulation of *Ggt1* mRNA and enzymatic activity compared to non-targeting controls (**Fig. 5E and F**). Ggt1 overexpression caused a minor defect in proliferation in cystine-containing medium, but fully restored cell numbers in cystine-free medium supplemented with GSH (**Fig. 5G and fig. S10A**).

We also observed a strong reduction in lipid peroxidation in Ggt1 overexpressing cells compared to controls in cystine-free medium supplemented with GSH (**fig. S10B and C**), confirming that extracellular GSH catabolism reduces oxidative stress in cystine deplete environments. Ggt1 overexpression also reduced the sensitivity to the GPX4 inhibitor RSL3 in the presence of cystine (**Fig. 5F**), suggesting that GSH recycling also contributes to ferroptosis protection.

To determine the effects of Ggt1 overexpression on GSH metabolism, we exposed the cells to different medium conditions for 6 hours. In cells exposed to cystine-free medium supplemented with GSH, Ggt1 overexpression strongly increased depletion of GSH from the medium and increased intracellular levels of cysteine, cystine and GSH (**fig. S10D and E**). Surprisingly, Ggt1 overexpression also increased intracellular GSH in cells after cystine starvation without GSH supplementation (**fig. S10E**), suggesting that Ggt1 participates in the recycling of GSH via the salvage pathway (*47*). We also noted that Ggt1 overexpression resulted in a small increase in cystine uptake from RPMI medium (**fig. S10D**), potentially also due to GSH recycling.

We next investigated the role of extracellular GSH catabolism in metastasis formation using a method developed to induce simultaneous establishment of lung, liver and kidney metastases (*48*)(**Fig. 5I**). Hydrodynamic tail vein injection of 5 x 10^5^ 4T1 cells into syngeneic mice resulted in efficient colonization of lungs and livers, as determined by histological analysis (**fig. S11A**). We therefore chose the lower number of 3 x 10^5^ cells to investigate the effect of Ggt1 overexpression on metastasis formation. Visual inspection of livers and lungs revealed clear metastatic nodules only in the livers of mice injected with Ggt1 overexpressing cells (**Fig. 5J**). Increased formation of liver metastases was also seen in histological sections while no metastatic nodules were observed in the lungs of the same animals (**Fig. 5K**). Quantification showed that Ggt1 significantly increased the number of metastatic nodules per liver section and the total metastatic area (**Fig. 5L**). These results demonstrate that the ability to degrade extracellular GSH increases the ability of cancer cells to colonize the cystine-poor microenvironment of the liver.

### Ggt1 drives spontaneous metastatic dissemination

Metastatic cancer cells are exposed to high levels of oxidative stress during invasion and dissemination via the vascular system (*11*). We hypothesized that catabolism of extracellular GSH could drive metastasis formation by protecting cancer cells under conditions of elevated oxidative stress also in environments where cystine is available at low concentrations (**Fig. 6A**). We therefore exposed control and Ggt1 overexpressing 4T1 breast cancer cells to plasma-like conditions, containing 30 µM cystine and 60 µM GSH. Elevated oxidative stress in context of ferroptosis was simulated by treatment with increasing doses of RSL3. As expected, Ggt1 overexpressing cells showed reduced RSL3 sensitivity under these conditions (**Fig. 6B and C**).

**Figure 6:**
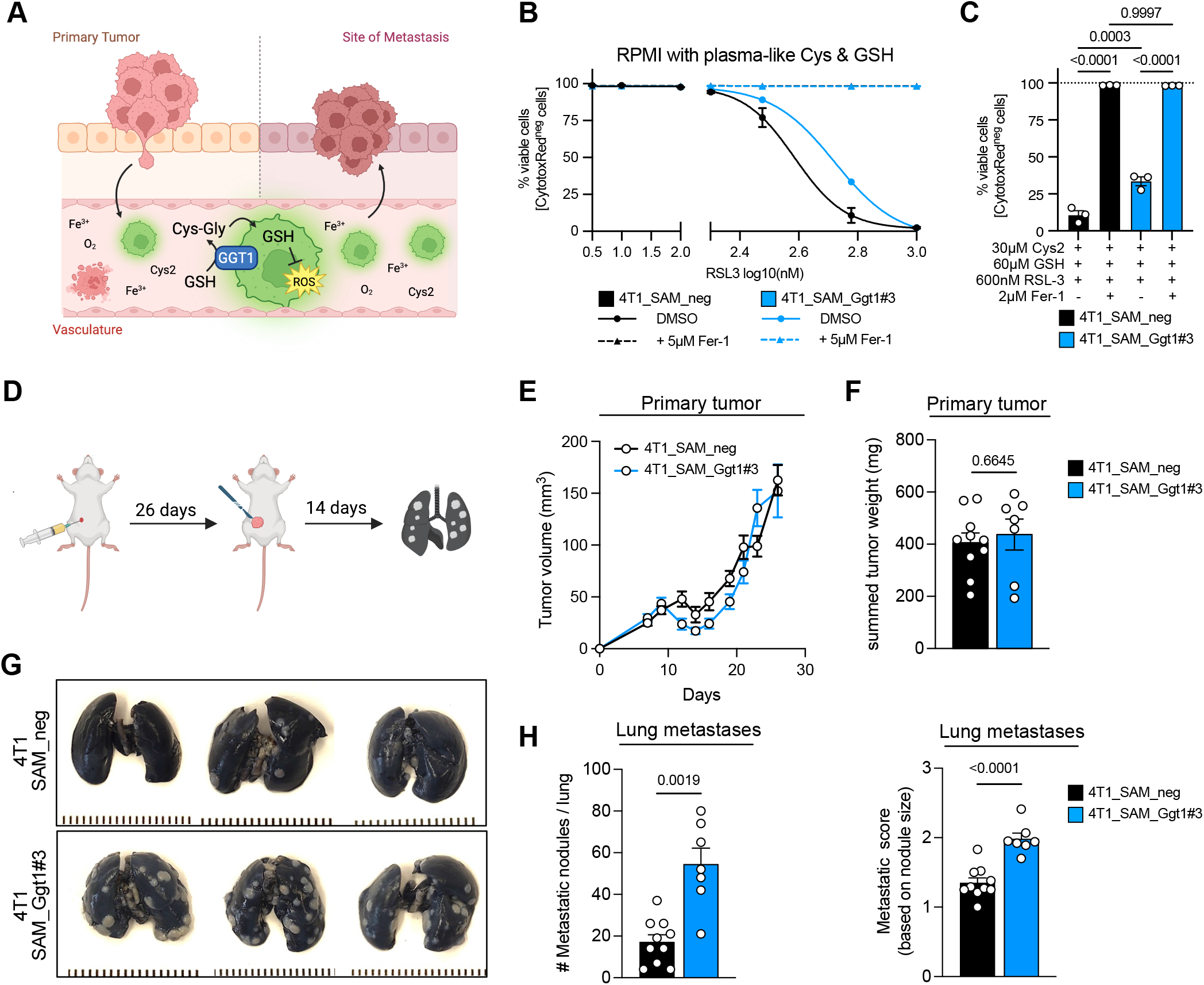
Ggt1 overexpression drives spontaneous metastasis in an orthotopic breast cancer model. (**A**) Diagram showing enhanced oxidative stress during invasion and dissemination of cancer cells via the vascular system. Created in https://BioRender.com. (**B**) 4T1 control and Ggt1 overexpressing cells were cultured under “plasma-like” conditions (30 µM cystine and 60 µM GSH) and treated with increasing doses of RSL3 either in the presence or absence of 4 µM ferrostatin (Fer-1). Viability was determined by automated cell imaging (Incucyte). Data show mean ± SD of biologically independent samples (n=4). (**C**) Comparison of viability in the presence of 600 nM RSL3. Significance was calculated using one-way ANOVA with Dunnett’s post-hoc test. (**D**) Schematic indicating spontaneous metastasis formation after orthotopic implantation into the 4^th^ mammary fat pad (MFP) followed by resection of the primary tumor after 26 days. Outgrowth of lung metastasis was evaluated after an additional 14 days. Created in https://BioRender.com. (**E**) Tumor volume of primary tumors determined by caliper measurements over 26 days. Data show mean ±SEM of individual tumors (n=20 for each group). (**F**) Summed tumor weight of resected primary tumors at 26 days from both flanks of each mouse. Data show mean ±SEM of individual tumors (4T1_SAM_neg, n=10; 4T1_SAM_Ggt1#3, n=7). Significance was calculated using unpaired t test with Welch’s correction. (**G**) Representative images of lungs from mice after orthotopic implantation and metastatic outgrowth of 4T1 control (SAM_neg) and Ggt1 overexpressing cells (SAM_Ggt1#3). Lungs were fixed and stained with India ink. (**H**) Quantification of metastatic nodules and metastatic score of lung metastatic lesions identified by India ink staining. Data represent mean ±SEM of individual lungs (4T1_SAM_neg, n=10; 4T1_SAM_Ggt1#3, n=7). Significance was calculated using unpaired t test with Welch’s correction.

We next investigated whether Ggt1 overexpression also affects spontaneous metastasis of primary tumors by applying an orthotopic implantation model that metastasizes to the lung. In this model, murine breast cancer cells (4T1) are implanted into the 4^th^ mammary fat pad. After initial growth for 26 days, the primary tumor was surgically removed, and mice were kept for a further 14 days to allow metastasis formation (**Fig. 6D**). While Ggt1 overexpression did not alter volume increase or weight at endpoint of primary tumors (**Fig. 6E and F**), it strongly increased the appearance of metastatic nodules in the lung (**Fig. 6G**). Quantification confirmed an increased number of metastatic nodules per lung and an elevated metastatic score (**Fig. 6H**). As this model has been optimized for the formation of lung metastasis (*49*), we did not detect any metastatic nodules in the liver (**fig. S11B**).

In conclusion, our study identified extracellular GSH catabolism as a mechanism that promotes proliferation and provides protection from oxidative damage, particularly under conditions of nutrient depletion and elevated stress. Targeting this mechanism could potentially provide an actionable benefit for cancer patients.

## Discussion

Therapeutic targeting of metastatic cancer remains a major clinical challenge. To identify selective vulnerabilities of metastasizing cancer cells, we determined the metabolic microenvironment of liver and lung, two organs frequently colonized by disseminated cancer cells (*3*). In accordance with previous studies (*50, 51*), we found that interstitial fluid (IF) from liver displayed a higher abundance of glutamine, isoleucine, leucine and methionine, while lung-IF contained higher levels of aspartate. In addition, cystine, the oxidized form of cysteine, was highly abundant in plasma and lung-IF but absent in liver-IF. In contrast, GSH was highly abundant in liver-IF, raising the possibility that cancer cells utilize extracellular GSH to compensate for the absence of cystine.

We found that extracellular GSH substitutes for environmental cystine in supporting cancer cell proliferation and preventing ferroptotic cell death in murine and human breast cancer and melanoma cell lines. Cell viability rescue required the activity of GGT, the enzyme responsible for extracellular GSH catabolism (*26*). Stable isotope tracing demonstrated that extracellular GSH catabolism provides substrates for *de novo* GSH synthesis. Viability rescue by extracellular GSH was independent of system Xc^-^, suggesting that cysteine derived from this process is maintained in its reduced form and taken up by other amino acid transporters, such as SLC4A1. Supplementation of culture medium with Cys-Gly, the direct product of the GGT reaction, was sufficient to restore cell viability and ferroptosis protection, but this was blocked by Xc^-^ inhibition, indicating that the lower reducing capacity of Cys-Gly is not sufficient to prevent cysteine oxidation. Both GSH and Cys-Gly restored intracellular GSH levels and prevented amino acid accumulation induced by cystine-starvation, indicative of active protein biosynthesis. Cys-Gly was more effective in blocking the formation of γ-glutamyl-dipeptides that has been shown to protect cancer cells viability by sequestering excess glutamate under cysteine-limiting conditions (*41*).

We found that GGT1 mRNA levels are associated with poor survival in several cancer entities and selectively upregulated in breast cancer liver metastases. Elevated expression of GGT1 and GCLC protein were also found in liver metastasis from breast cancer patients with advanced and treatment-refractory disease. Expression of GGT enzymes is induced in response to oxidative stress via the Ras/NRF2 and NF-κB axis (*52–54*), suggesting that elevated oxidative stress during metastasis formation drives GGT1 expression to increase the anti-oxidant capacity of cancer cells. We observed that overexpression of Ggt1 in murine breast cancer cells provided protection from lipid peroxidation and ferroptosis in cystine-low conditions, confirming the role of extracellular GSH catabolism in cellular redox homeostasis. Furthermore, overexpression of Ggt1 increased the ability of murine breast cancer cells to produce liver metastases after hydrodynamic tail vein injection in syngeneic mice. This result is in line with a previous study indicating that GGT overexpression promotes liver metastasis in B16-F1 melanoma cells after intrasplenic injection (*55*). In addition, Ggt1 overexpression increased spontaneous metastasis formation of breast cancer cells in an orthotopic implantation model. This suggests that GSH catabolism protects cancer cells from oxidative stress imposed during the different stages of metastatic spread, such as detachment from the extracellular matrix, migration and invasion, and dissemination via the vascular system.

Our results are in line with recent findings demonstrating that catabolism of extracellular GSH provides cancer cells with an alternative source of cysteine (*29*). We now extend this work by showing that catabolism of extracellular GSH enables cancer cells to selectively colonize cystine-deprived environments, such as the liver. In contrast to the study by Hecht and colleagues, we did not observe any effect of Ggt1 overexpression on primary tumor growth in our syngeneic breast cancer model, indicating site-specific differences or effects mediated by immune interactions. Indeed, low-level oxidative stress can be pro-tumorigenic by promoting immune evasion (*56*), encouraging further studies. Together, our results demonstrate that catabolism of extracellular GSH supports cancer cell survival in metabolically deprived microenvironments and under conditions of elevated oxidative stress, thus identifying GGT as potential therapeutic target.

## Supporting information

Vogel et al_Table S1_patient information

Vogel et al_Data S1

## Acknowledgments

We would like to thank S. Haid, I. Germer and I. Koltsaki for technical assistance. We also thank Stephanie Laier, Hannah Walzer and Julia Förderer from the Core Unit Tumor Models for help with animal experiments and Thomas Metzler from the Core Unit Preclinical and Translational Pathology for input on histological analysis of mouse tissue. We are grateful to the DKFZ Core Unit Light Microscopy for their support.

## Funding

This work was funded by the following grants:

German Research Foundation grant SPP-2306, SCHU2670/2-1/2 (FCEV, DS, JH, AS)

German Research Foundation grant FOR-2314, SCHU2670/2-1/2 (FR, MTS, AS)

German Research Foundation grant SCHU2670/1-2 (FR, MTS, AS)

Excellence through Focus’ funded by zukunft.niedersachsen, the joint science funding program of the Lower Saxony Ministry of Science and Culture and the Volkswagen Foundation, UMG2032 (FCEV)

German Academic Exchange Service (KMA)

This study was supported in part by the National Center for Tumor Diseases (NCT), Heidelberg (CATCH)

The MSNZ Würzburg is funded by the German Cancer Aid, NG2 (AR)

IZKF Würzburg grant C-524 (AR)

Else Kröner-Fresenius-Stiftung grant 2022_EKEA.76 (AR)

## Author contributions

Conceptualization: FCEV, FR, AR, AS

Methodology: FCEV, ABCF, LS

Investigation: FCEV, FR, DS, JH, AMW, HR, MTS, KMA, PP, VF, TP

Data curation: FCEV, ABCF, LS, CS, BK

Funding acquisition: FCEV, KMA, AS

Resources: LW, VT, AS*, PL

Writing – original draft: FCEV, AS

Writing – review & editing: all authors

Supervision: AS

*Andreas Schneeweiss

## Data, code, and materials availability

All data are available in the manuscript or the supplementary materials. All unique and stable reagents generated in this study are available from the lead contacts on reasonable request with a completed materials transfer agreement.

## Supplementary Materials

Materials and Methods

Figs. S1 to S11

Tables S1 to S3

Data S1

## Materials and Methods

### Chemicals and Inhibitors

Ferrostatin-1 (#17729), Liproxstatin-1 (#17730) and BSO (#14484) were obtained from Cayman Chemicals Corp. (Ann Arbor, MI). (1S,3R)-RSL3 (#6118), Z-VAD-FMK (#2163) and GGsTOP (#4452) were purchased from Bio-Techne / Tocris. Erastin (#329600), L-Cysteine-L-Glycine (Cys-Gly; #C0166), and Glycylglycin (Gly-Gly; #G3915) were purchased from Sigma-Aldrich / Merck. Glutathione (GSH; #6832.2) was purchased from Carl Roth. BODIPY™ 581/591 C11 (#D3861) and β-Mercaptoethanol (#31350010) were purchased from Thermo Fisher Scientific. Incucyte® CytotoxRed (#4632) or CytotoxGreen (#4633) Dye for Counting Dead Cells were purchased from Sartorius.

### Cell culture

4T1 cells were from American Tissue Type Collection (ATCC, CRL-2539. E0771 cells were kindly provided by F. Loayza-Puch (DKFZ, Heidelberg, Germany). B16-F1 and A375 were kindly provided by A. Tasdogan (University Hospital, Essen, Germany). MDA-MD-231, MDA-MD-436 and MDA-MD-468 were originally obtained from LRI Cell Services (CRUK, LRI, London, UK). All cell lines were used at low passage and frequently checked for absence of mycoplasma. Murine cell lines were cultured in RPMI (Sigma Aldrich #R0883) with 10% fetal bovine serum (FBS), 2 mM L-glutamine, 100 U/mL penicillin, and 100 μg/mL streptomycin. Human cell lines were cultured in DMEM (Gibco #21969-035) with 10% FBS, 2 mM L-glutamine, 100 U/mL penicillin, and 100 μg/mL streptomycin (all Sigma). Cells were passaged in medium containing standard FBS. Medium containing dialyzed FBS was used for all experiments. Dialysis of FBS was performed using SnakeSkin (MWCO 3.5kDa, Thermo Fisher #88244) against 10x volume of saline solution at 4 °C before sterile filtration. RPMI (Sigma Aldrich #R7513) and DMEM (Sigma Aldrich #D0422) culture medium without cystine was used with or without the addition of glutathione (Carl Roth #6832.2); in this medium L-methionine was added back to match concentration of corresponding normal RPMI and DMEM culture medium. All cell lines were cultured at 37 °C in a humidified incubator at 5% CO_2_.

### Genetic modification using CRISPR

For CRISPR/Cas9 mediated deletion or overexpression of murine *Ggt1*, sgRNA sequences were designed using CHOPCHOP (https://chopchop.cbu.uib.no), VBC-score (https://www.vbc-score.org), or Broad CRISPick tool (https://portals.broadinstitute.org/gppx/crispick/public). Cloning for knockout was done into *lentiCRISPRv2* (Addgene #52961) and for overexpression into *lentiSAMv2* (Addgene #75112). For overexpression target cells were transduced with the helper plasmid *lentiMPH v2* (Addgene #89308) prior to transduction with *lentiSAMv2*. Viral particles were produced in HEK293T cells and stable populations were selected using respective agent and used as pools. Silencing and knockout efficiency was confirmed using RT-qPCR. lentiCRISPR v2 (Addgene #52961), lentiSAMv2 (Addgene #75112), and lentiMPH v2 were a gift from Feng Zhang (*57, 58*). Primers are provided in Table S2.

### Gene expression analysis with RT-qPCR

RNA was extracted using TRI-Reagenz (Sigma Aldrich #T9424) following the standard manufactures protocol. 2 μg of each RNA sample were reverse transcribed using M-MLV RT (Promega #M1701) following manufactures protocol. Real-time qPCR was performed using PowerUp SYBR Green Master Mix (Applied Biosystems #A25777) on a StepOne real-time PCR instrument (Applied Biosystems). The relative expression quantity was normalized to housekeeper gene *Ppia* (murine) or *GUSB* (human). Relative expression was normalized against the control group where indicated. Primers are provided in Table S3.

### GGT activity assay

Enzymatic activity of γ-Glutamyltransferase (GGT) were analyzed by colorimetric Activity Assay (Sigma Aldrich #MAK089) following the standard manufactures protocol. Briefly, around 1*10^6 cells or 10µl RPMI and dFBS samples were lysed and homogenized with 200µl ice-cold GGT Assay Buffer and centrifuged to remove insoluble material. GGT activity is determined by transfers of the γ-glutamyl group from the substrate L-γ-Glutamyl-*p*-nitroanilide, liberating the chromogen p-nitroanilide (*p*NA, 418 nm) proportional to the GGT present; a parallel *p*NA Standard curve is applied. Absorbance was measured at 418 nm in a kinetic every 3 min at 37 °C in the plate reader. GGT activity data from cell lysates are finally normalized to protein content of the lysate.

### Analysis of cell viability and proliferation using crystal violet

Cells were seeded in 24 or 96 well plates and treated as indicated. After incubation, cells were washed with PBS and fixed for 10 minutes in 10% buffered formalin (Carl Roth #P087.4). Cells were washed and stained with 0.1% crystal violet (Sigma #C0775) solution containing 20% EtOH for 30 min. Plates were carefully rinsed in tap water, dried, scanned and extracted using 10% acetic acid. Absorbance was measured at 550 nm.

### Live cell imaging of cell viability and lipid peroxidation

Real-time monitoring of cell proliferation, cell viability and lipid peroxidation was conducted using the IncuCyte® S3 Live-Cell Analysis System (Sartorius). Briefly, cells were seeded in 24-well or 96-well plates and allowed to adhere overnight prior to the experiments. For viability assessments, cultures were treated with IncuCyte® CytotoxRed or CytotoxGreen dye (Sartorius) to identify dead cells, following the manufacturer’s protocols. Brightfield and fluorescent images were captured in different intervals (between every 1 to every 8 hour) using the IncuCyte system sitting in a humidified incubator at 37 °C and 5% CO_2_. Quantification of live and dead cells was also performed using the integrated IncuCyte AI Cell Health Analysis Software Module.

To assess lipid peroxidation in live cells seeded in a 24-well format, we used the C11-BODIPY 581/591 sensor (Thermo Fisher Scientific #D3861). Briefly, after washing once with prewarmed HBSS+/+ buffer (Gibco #14025-092), cells were incubated with 5 µM C11-BODIPY 581/591 in HBSS+/+ at 37°C for 20 minutes. After this, cells were washed again with HBSS+/+ buffer and treatment conditions were added to the wells in full RPMI or DMEM medium. Cells were then imaged with brightfield, red, and green fluorescence channels of the IncuCyte system allowing analysis of lipid peroxidation levels over time and on live cells (images were captured in intervals of 1 or 2 hour). Image analysis was conducted using the AI Cell Health Analysis Software Module and subsequent Cell-By-Cell Analysis Software Module to quantify the dynamic changes in lipid peroxidation positive cells.

### Isolation of interstitial fluids

For the harvesting of organ interstitial fluids of healthy wildtype mice, 7-8 weeks old female Balb/c were used. Isolation of the organ interstitial fluids (IF) was performed with the full organ by careful centrifugation on a nylon mesh taped onto a 50ml falcon tube as previously described (*59*). From the same animals, blood was isolated from the heart by cardiac puncture and plasma was enriched by subsequent centrifugation for 10min with 840g at 4°C in EDTA-coated plasma collection tubes (Sarstedt #41.1395.105). Plasma and IF were snap frozen and stored at -80°C until further processing.

### LC-MS Metabolomics

For intracellular metabolites, cells were washed with cold saline solution (B.Braun # 3570130), snap frozen in liquid nitrogen and scraped off in 500µl ice-cold MeOH/H_2_O/Acetonitrile (50/20/30 v/v) containing internal standards (MSK-CAA-1, DLM-7654, DLM-9476, DLM-9045, DLM-9071, DLM-6068, DLM-831, DLM-3487 Cambridge Isotope laboratories). 50µl of each media supernatant sample was mixed with 450µl of MeOH/H_2_O/Acetonitrile (50/20/30 v/v) containing all standards. For metabolomics of thiol-containing molecules, NEM-derivatization protocol was applied as published (*36, 37*).

Metabolite extraction was performed by solid phase extraction (SPE) following standard protocols. Dried metabolite extracts were dissolved in 100 μl 5 mM NH_4_OAc in CH_3_CN/H_2_O (75:25, v/v) or 100 μl CH_3_CN/H_2_O (75:25, v/v), and 3 µl of each sample was applied to an amide-HILIC column (2.6 μm, 2.1×100 mm, Thermo Fisher, 16726–012105). Metabolites were separated at 30 °C by LC using a DIONEX Ultimate 3000 UPLC system and the following solvents: solvent A consisting of 5 mM NH_4_OAc in CH_3_CN/H_2_O (5:95, v/v) and solvent B consisting of 5 mM NH_4_OAc in CH_3_CN/H_2_O (95:5, v/v). The LC gradient program was: 98% solvent B for 2 min, followed by a linear decrease to 40% solvent B within 5 min, then maintaining 40% solvent B for 13 min, then returning to 98% solvent B in 1 min and then maintaining 98% solvent B for 5 min for column equilibration before each injection. The flow rate was maintained at 350 μl/min. The eluent was directed to the hESI source of the Q Exactive mass spectrometer (QE-MS; Thermo Fisher Scientific) from 1.85 min to 18.0 min after sample injection. The scan range was set to 69.0–1000 m/z with a resolution of 70,000 and polarity switching (negative and positive ionization). Peaks corresponding to the calculated metabolites masses taken from an in-house metabolite library (MIM +/− H+ ± 2 mmU) were integrated using the El-MAVEN software v.12.1-beta (*60*).

### Absolute quantification of liver, lung and plasma metabolites

For absolute metabolite quantification of liver-IF, lung-IF and plasma, we designed a standard addition approach using fetal bovine serum (FBS) as a surrogate complex matrix with unlimited availability and high reproducibility. Different volumes of FBS (2 or 5 µl) were spiked with either 25%, 50% or 100% volume of mouse plasma like medium (MPLM), formulated with defined metabolite concentrations (*32, 33*). As MPLM does not contain GSH, this metabolite was quantified separately using standard dilutions in FCS at defined concentrations. Signal intensities obtained by LC-MS were plotted against spiked concentrations to determine the respective metabolite concentrations in FCS from the x-intersect of the regression line. Fixed volumes of fluids (5 µl for liver-IF; 2 µl for lung-IF; 10 µl for plasma) were extracted and analyzed by LC-MS. Absolute metabolite concentrations were calculated using the same volume of FBS extracted and analyzed in parallel as standard. Lactate levels were quantified using the Lactate-Glo™ Assay kit (Promega, #J5021) following the manufacturers protocol.

### Stable isotope tracing

Stable isotope labelled metabolites for GSH (glycine-^13^C_2_^15^N_1_; #CNLM-6245-HP-10) or L-Glutamine (^13^C_5_; #CNLM-1822-H-0.1) were purchased from Cambridge Isotope Laboratories. Isotopologue peaks were integrated using the El-MAVEN software and the R package IsoCorrectoR (*61*) was used to correct for natural isotope abundance.

### Analysis of public gene expression and survival data

We selected gene expression dataset *GSE110590* containing data from human samples of breast cancer metastases (*45*). Data were downloaded and used with the already processed and normalized data matrix. Gene set enrichment analyses (GSEA) were performed using MSigDB v5.2 with default parameters and 1000 permutations using either the KEGG-LEGACY or the WikiPathway subset of canonical pathways within the C2 gene sets.

Kaplan-Meyer curves of TCGA breast cancer (BRCA), Pancreatic adenocarcinoma (PDAC) and Lung squamous carcinoma (LUSC) were generated using the human protein atlas (https://www.proteinatlas.org). Box Plot stage dependent analysis of GGT1 expression of TCGA Breast cancer data (BRCA) were performed using GEPIA2 (*62*). Correlation of gene expression and cancer dependencies with small molecules was done with the BROAD cancer therapeutics response Portal v2 (https://portals.broadinstitute.org/ctrp.v2/).

### Patient biomaterial and ethics approval statement

The collection of human breast cancer samples (primary and metastasis) was derived from patients participating in the CATCH (Comprehensive Assessment of Clinical Features and Biomarkers to Identify Patients with Advanced or Metastatic Breast Cancer for Marker Driven Trials in Humans) and COGNITION (Comprehensive assessment of clinical features, genomics and further molecular markers to identify patients with early breast cancer for enrolment on marker driven trials) trial at the Division of Gynecologic Oncology, NCT Heidelberg. Written informed consent was retrieved according to the study approval protocol by the local IRB of Heidelberg University (S-164/2017 (NCT05652569) and S-790/2018 (NCT05906407)).

### Proteomics of patient biomaterial

Prior to sample preparation, frozen tissue sections were assessed by pathologists at the NCT Heidelberg to determine vital tumor cell content via H&E staining according to a standard protocol. Information on patients and prior treatment is provided in Table S1.

Cell lysates from fresh-frozen tumor specimens were obtained using the AllPrep DNA/RNA/miRNA Universal Kit and the QIAamp DNA Mini Kit (Qiagen) at the NCT Heidelberg Sample Processing Laboratory. Cell lysate protein concentrations were determined using the Pierce Bradford Protein Assay Kit (Thermo Fisher). Preparation of samples for mass-spectrometric analysis was performed in batches of nine patient samples and two reference samples (200 µg per sample) per tandem mass tag (TMT) batch as previously described with minor adaptations (*63*). For LC/MS Proteomics data acquisition, multiplexed samples were measured on an Orbitrap Fusion Lumos Tribrid Mass Spectrometer (Thermo Fisher) coupled to Vanquish Micro-Flow HPLC system (Thermo Fisher) for whole-proteome analysis. For peptide identification and quantification, raw files were searched with MaxQuant (version 2.1.3.0) against the human UniProt database, including unreviewed proteins (97,086 sequences including isoforms) and the MaxQuant common contaminants database. Identifications were transferred with SIMSI-Transfer to improve data completeness across TMT batches (*64*). Median centering data transformation was performed to normalize data within- and across-batch. Protein groups were inferred and quantified at the gene level using the Picked Protein Group FDR package (*65*). Of note, data for GGT1 include integrated data from pseudogene GGT2 with identical peptide sequence.

### Ethics approval of animal experiments

All animal experiments have been approved by committees of the regional authorities of the state of Baden-Württemberg (Regierungspräsidium Karlsruhe, G-200/22) or Bavaria (Regierung von Unterfranken, RUF-55.2.2-2532-2-2207). All mice were housed and maintained under pathogen free conditions with a 12-h light–dark cycle, a 45–65% relative humidity, temperature of 22 +/- 2 °C and ad libitum access to water and food in accordance with the institutional guidelines.

### Hydrodynamic tail vein injections

Hydrodynamic tail vein injection of murine cancer cells was performed in 6-8 week old female Balb/c wildtype mice. Briefly, mice were randomly allocated to the different groups and injected with either 3×10^5^ or 5×10^5^ 4T1 cells in a 0.9% NaCl solution to a final volume of 9% of mouse body weight within 5-8 seconds as described previously (*48*). Animals that were not efficiently injected within 8 seconds were excluded from the study. Mice were sacrificed after 14 days and tissues were harvested for subsequent analysis.

### Orthotopic implantation

To induce mammary tumors, 6-8 week old female Balb/c wildtype mice were orthotopically injected bilaterally into the fourth mammary fat pad. Each fat pad was injected with 7,5 x 10^5^ 4T1-SAM_neg or 4T1-SAM_Ggt1#3 cells resuspended in 50 µL PBS. Tumor size was measured with calipers, and the volume was calculated using the ellipsoid formula π/6 × (length × width^2^). Primary tumors were surgically resected after 26 days; their size and weight were measured. At endpoint, 14 days after primary tumor resection, animals were euthanized, weighed, and lungs and livers were collected. Animals were excluded from the study if tumors failed to form, tumors were growing in the peritoneum, or if health concerns were reported.

### Histological analysis

After overnight fixation in 10 % buffered formalin, representative tissue specimens of murine lungs and livers were dehydrated, embedded in paraffin, and cut into 4 μm-thick sections. Tissue sections were stained with H&E according to a standard protocol. H&E-stained slides were scanned at 20x resolution using a slide scanner (Zeiss Axioscan 7). Liver tumor nodules were analyzed using Fiji ImageJ (version 2.16.0) for individual size and quantification. Additional pathological assessment was performed on lung and liver sections without distinct nodules to confirm absence of metastases.

### Detection of lung metastasis after orthotopic implantation

India ink was used to visualize macroscopic 4T1 lung metastatic nodules as previously described (*66*). Briefly, 3 ml of India ink (15% black ink in PBS, Angene) was injected into the exposed trachea of sacrificed mice before the lungs were dissected. Lungs were then fixed in de-staining solution (3.7% formaldehyde (Thermo Fisher Scientific), 1.7% acetic acid (Carl Roth) in 70% ethanol (Sigma Aldrich)) overnight. Nodules were counted by two blinded observers and their size categorized as small (<1×1 mm, score 1), intermediate (1×1-2×2 mm, score 2), and large (>2×2 mm, score 3) under a dissecting microscope (Olympus). The metastatic score was calculated as (sum of the individual scores for each nodule)/(the total number of nodules), such that the score for each lung ranged from 1 to 3, depending on the average nodule size.

### Statistical analysis

Statistical analyses of data obtained from metabolomics were performed with MetaboAnalyst version 6.0 (www.metaboanalyst.ca). Statistical analyses of the other data were carried out with GraphPad Prism 10. Groups were compared by multivariate (PCA) and/or univariate analyses (t-test or ANOVA, p < 0.05, FDR-applied) followed by post-hoc tests unless otherwise stated in the figure legend. No statistical methods were used to pre-determine sample sizes for cell culture experiments. Power analysis was performed to determine size of animal cohorts based on pilot experiments (type 1 error = 5%; type 2 error = 20%; Software R, Version 4.3.0). It is stated in figure legend when data were log2 transformed before statistical analysis. Randomization was performed for animal experiments. Image analysis of tissue samples was performed in a double blinded manner for liver sections and using automated script for lung sections (Fiji ImageJ script for analysis of lung metastasis will be made available upon request). No animals or data points were excluded. All data were presented as average ± SD of independent biological replicates with number of replicates stated in figure legend.

---

**Figure S1:**
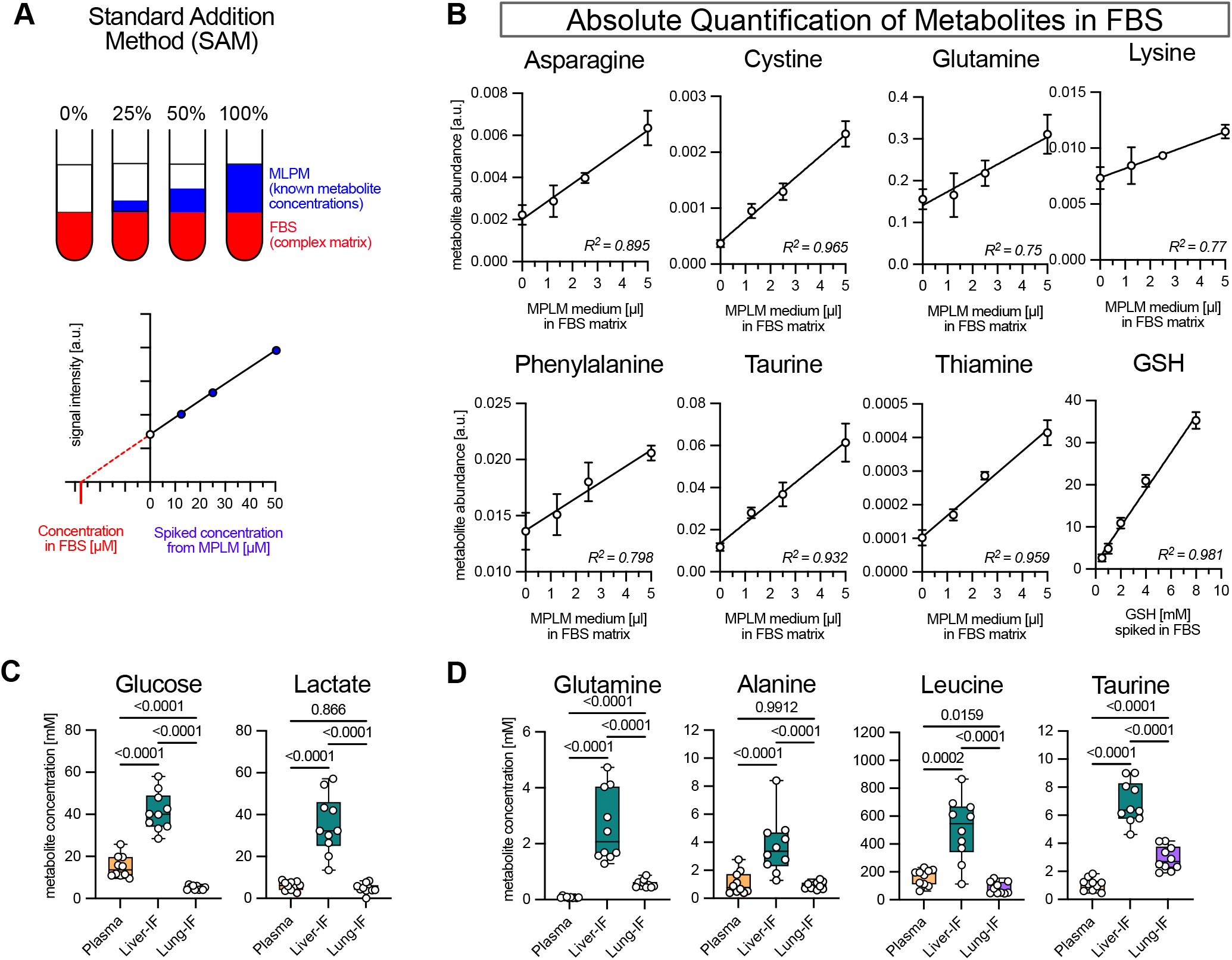
Absolute quantification of organ-specific metabolic microenvironments by Standard Addition Method (SAM). **(A)** Schematic overview of the standard addition method (SAM) for the absolute metabolite quantification in FBS as a complex biofluid with a comparable matrix to the murine plasma and organ interstitial fluids. MPLM = Mouse Plasma-Like Medium. **(B)** Representative graphs confirming linearity between signal intensity and cumulative standard addition using MPLM with known metabolite concentrations. Metabolite concentrations in FBS were determined from the intersection between the regression line and the x-axis. As GSH was absent from MPLM, a dedicated calibration curve was used. R-square values represent the Goodness of Fit of simple linear regression analysis. **(C)** Absolute metabolite concentrations of glucose and lactate. L-lactate concentrations were measured by Lactate-Glo Assay. Data are displayed as box and whisker plots, showing mean, 25th and 75th percentile and min-max. Significance was calculated using one-way ANOVA with Tukey post-hoc test on log transformed data (n=10). **(D)** Metabolite concentrations of selected amino acids (glutamine, alanine, leucine, and taurine) showing enrichment in liver-IF. Data are displayed as box and whisker plots, showing mean, 25th and 75th percentile and min-max. Significance was calculated using one-way ANOVA with Tukey post-hoc test on log transformed data (n=10).

**Figure S2:**
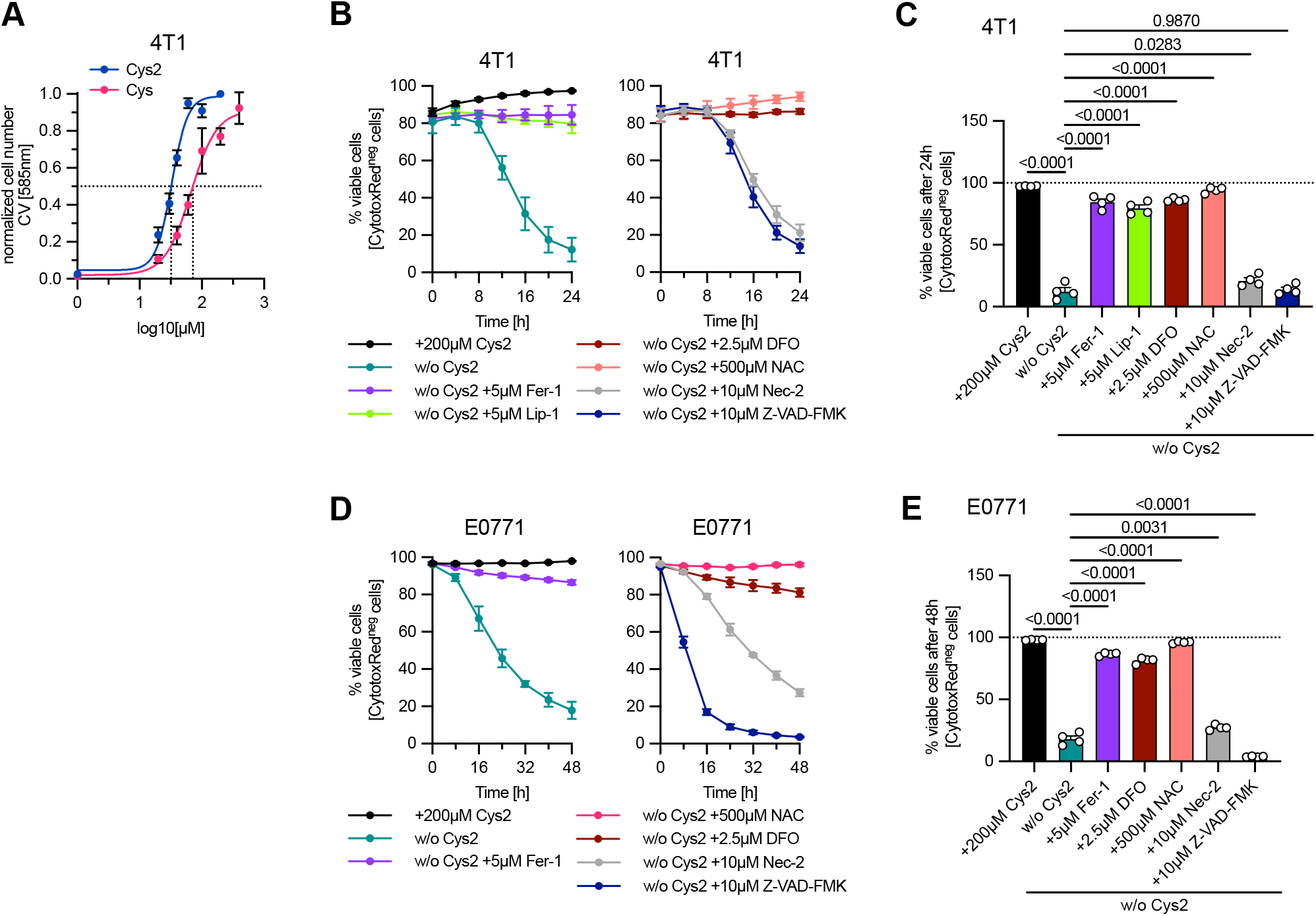
Cystine provision is essential for the viability and proliferation of murine breast cancer cells and is rescued by inhibition of ferroptosis. **(A)** 4T1 cells were cultured in RPMI medium containing different concentrations of cysteine (Cys) or cystine (Cys2) for 72 hours. Relative cell numbers were determined by crystal violet staining. Lines indicate IC50 for cysteine (66.6 µM) and cystine (33.0 µM). Data show mean ± SD of biologically independent samples (n=4). **(B)** Cell viability of 4T1 cells cultured under the indicated conditions determined by CytotoxRed staining and live cell imaging over time. **(C)** Cell viability data of 4T1 cells for 24h as shown in fig. 2B. Data show mean ± SD of biologically independent samples (n=4). Significance was calculated using one-way ANOVA with Dunnet post-hoc test. **(D)** Cell viability of E0771 cells cultured under the indicated conditions determined by CytotoxRed staining and live cell imaging over time. **(E)** Cell viability data of E0771 cells for 24h as shown in fig. 2B. Data show mean ± SD of biologically independent samples (n=4). Significance was calculated using one-way ANOVA with Dunnet post-hoc test.

**Figure S3:**
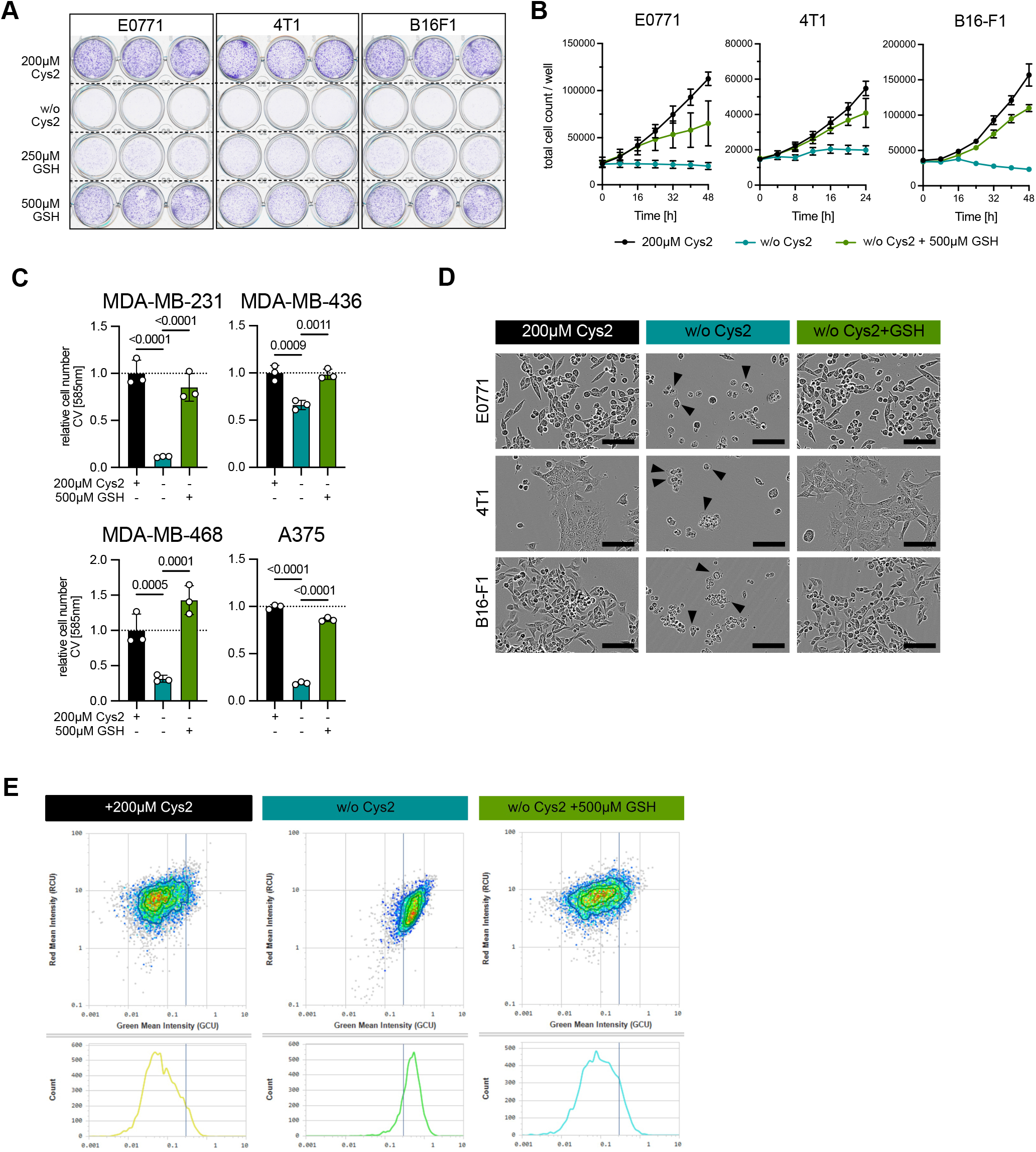
Extracellular GSH supports viability of murine and human breast cancer and melanoma cells and prevents lipid peroxidation. **(A)** E0771, 4T1and B16-F1 cells were cultured in RPMI containing 200 µM cystine (Cys2) or in cystine-free medium with or without addition of 250 or 500 µM GSH. Cells were cultured for 24 (4T1) or 48 (E0771 and B16F1) hours and stained with crystal violet (n=3). **(B)** Total cell counts of E0771, 4T1 and B16-F1 cells cultured in RPMI containing 200 µM cystine (Cys2) or in cystine-free medium with or without addition of 500 µM GSH for the indicated times determined by live cell imaging. Values represent means ±SD of biologically independent samples (n=4). **(C)** Human Breast cancer cells (MDA-MB-231, MDA-MD-436 and MDA-MB-468) and human melanoma cells (A375) were cultured under the indicated conditions for 72 hours. Relative cell number was determined by crystal violet staining normalized to respective controls. Data show mean ± SD of biologically independent samples (n=3). Significance was calculated using one-way ANOVA with Tukey post-hoc test on log transformed data. **(D)** Representative images of E0771, 4T1 and B16-F1 cells cultured in the indicated conditions for 24 hours. Arrowheads indicate cells displaying cytoplasmic swelling and leakage as characteristics of ferroptosis. Scale bar 200µm. **(E)** Representative scatter plots and histograms indicating the shift in BODIPY_581/591_-C_11_ green fluorescence in 4T1 cells cultured in the absence of cystine (w/o Cys2) and its restoration by addition of GSH after 20 hours analyzed by live cell imaging.

**Figure S4:**
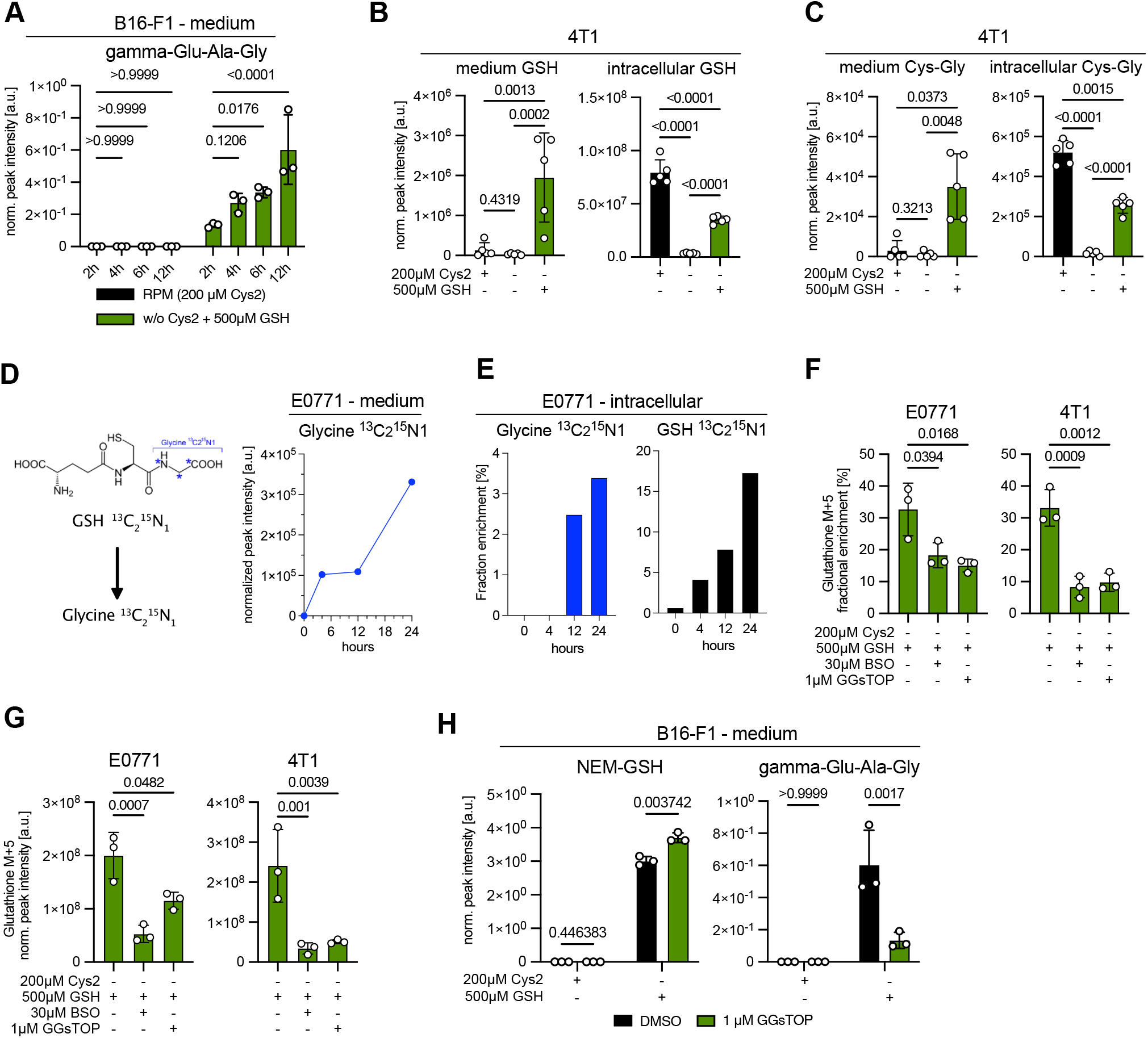
Cancer cells degrade extracellular GSH using a gamma-glutamyl transferase. **(A)** B16-F1 cells were incubated in RPMI containing 200 cystine (Cys2) or cystine-free medium supplemented with 500 µM GSH for 12 hours. At the indicated times, medium samples were taken and analyzed for the appearance of γ-glutamyl peptides produced by the transpeptidase activity of GGT enzymes. Graph shows relative abundance of γ-glutamyl-alanyl-glycine (γ-Glu-Ala-Gly). Data show mean ± SD of biologically independent samples (n=3). Significance was calculated using two-way ANOVA with Dunnet post-hoc test. **(B)** 4T1 cells were cultured in RPMI containing 200 µM cystine (Cys2) or in cystine-free medium with or without addition of GSH for 6 hours. Relative metabolite abundance of GSH in medium and cell extracts was determined by LC-MS. Data show mean ± SD of biologically independent samples (n=5). Significance was calculated using one-way ANOVA with Tukey post-hoc test on log transformed data. **(C)** Relative metabolite abundance of cysteinyl-glycine (Cys-Gly) in medium and cell extracts as in fig. S4B. **(D)** Schematic overview of the stable isotope labelling using GSH-^13^C_2_^15^N_1_ (left). E0771 cells were cultured in cystine-free medium containing 500 µM GSH-^13^C_2_^15^N_1_. After the indicated times, medium samples were extracted and analyzed for the abundance of Glycine-^13^C_2_^15^N_1_ (right). Data represent single replicates across multiple time points. **(E)** Fractional enrichment of intracellular Glycine-^13^C_2_^15^N_1_ and GSH-^13^C_2_^15^N_1_ of E0771 cells cultured in cystine-free medium containing 500 µM GSH-^13^C_2_^15^N_1_. Data represent single replicates across multiple time points. **(F)** E0771 and 4T1 cells were pre-labelled in standard RPMI medium supplemented with L-Glutamine-^13^C_5_ for 24 hours. After this time, cells were switched to indicated culture condition and treated with 30 µM buthionine sulphoximine (BSO) or 1 µM GGsTOP in cystine-free medium with addition of 500 µM GSH, still in presence of L-Glutamine-^13^C_5_. Samples were taken after 24 hours and fractional enrichment of GSH M+5 in cells from was quantified by LC-MS. Data show mean ± SD of biologically independent samples (n=3). Significance was calculated using one-way ANOVA with Dunnett post-hoc test. **(G)** Relative metabolite abundance of GSH M+5 in E0771 and 4T1 cell extracts as in fig. S4F. **(H)** B16-F1 cells were incubated in RPMI containing 200 cystine (Cys2) or cystine-free medium supplemented with 500 µM GSH with or without 1µM GGsTOP for 12 hours. Medium samples were taken and analyzed for the presence of GSH (NEM-GSH) and appearance of γ-glutamyl-alanyl-glycine (gamma-Glu-Ala-Gly). Data show mean ± SD of biologically independent samples (n=3). *NB* Data points for DMSO control conditions are also shown in fig. S4A. Significance was calculated using two-way ANOVA with Bonferroni post-hoc test.

**Figure S5:**
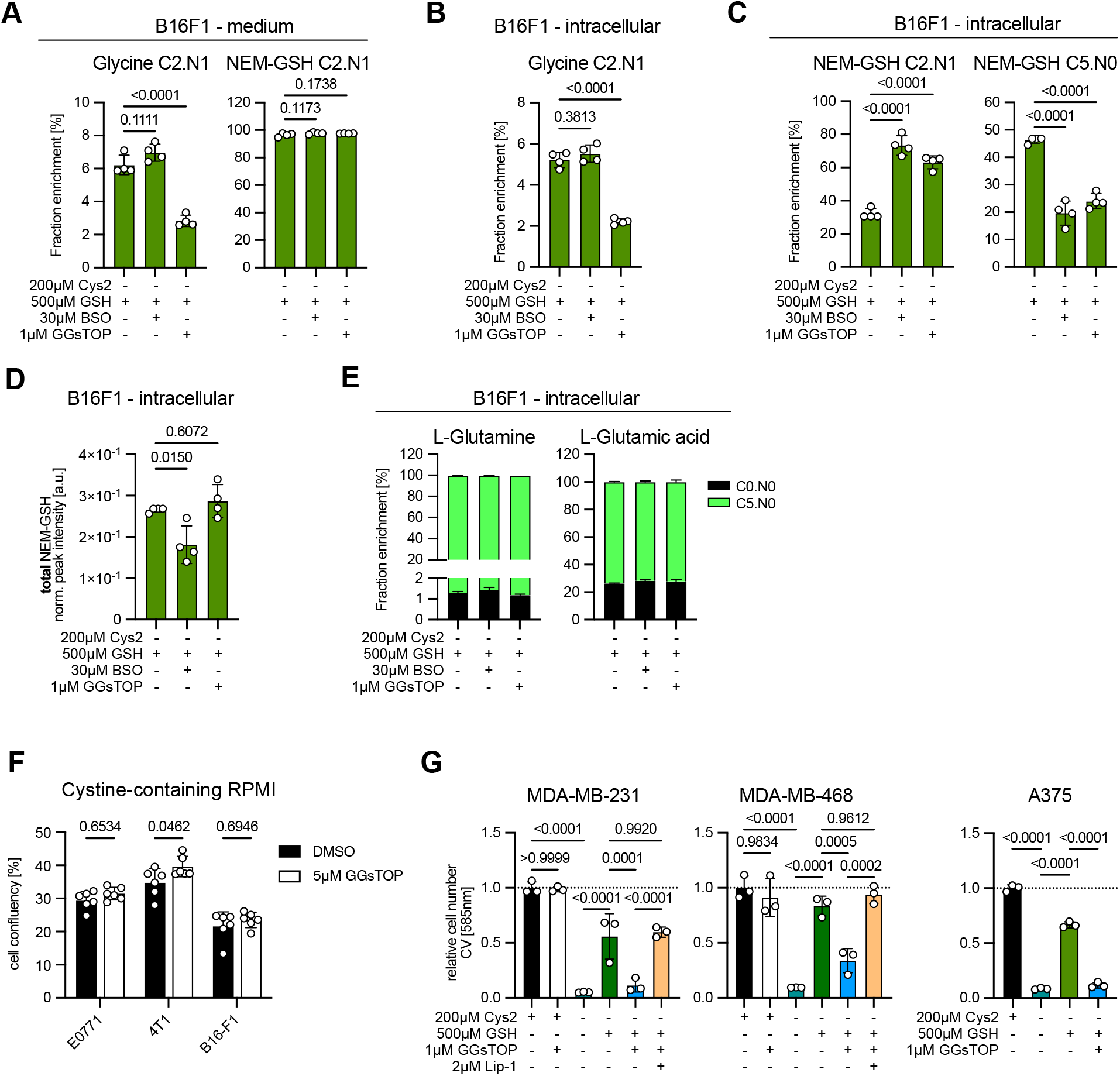
Cancer cells take up intact GSH and degrade extracellular GSH via GGT and also to generate substrates for *de novo* GSH synthesis. **(A)** B16F1 cells were pre-labelled in standard RPMI medium supplemented with L-Glutamine-^13^C_5_ for 24 hours. After this time, cells were switched to indicated culture condition and treated with 30 µM BSO or 1 µM GGsTOP in cystine-free medium with addition of GSH-^13^C_2_^15^N_1_ and L-glutamine-^13^C_5_. Samples were taken after 24 hours and fractional enrichment of Glycine-C2.N1 and NEM-GSH C2.N1 in medium was quantified by LC-MS thiol metabolomics. Data show mean ± SD of biologically independent samples (n=4). Significance was calculated using one-way ANOVA with Dunnett post-hoc test. **(B)** Fractional enrichment of Glycine-C2.N1 in B16F1 cell extracts as in fig. S5A. Data show mean ± SD of biologically independent samples (n=4). Significance was calculated using one-way ANOVA with Dunnett post-hoc test. **(C)** Fractional enrichment of NEM-GSH C2.N1 and NEM-GSH C5.N0 in B16F1 cell extracts as in fig. S5A. Data show mean ± SD of biologically independent samples (n=4). Significance was calculated using one-way ANOVA with Dunnett post-hoc test. **(D)** Total intracellular summed metabolite abundance of NEM-GSH in B16F1 cell extracts as in fig. S5A. Data show mean ± SD of biologically independent samples (n=4). Significance was calculated using one-way ANOVA with Dunnett post-hoc test. **(E)** Fractional labelling of intracellular L-Glutamine and glutamic acid in B16F1 cell extracts as in fig. S5A. **(F)** E0771, 4T1 and B16-F1 cells were cultured in RPMI and treated with 5µM GGsTOP for 24 hours. Confluency was determined by live cell imaging. Data show mean ± SD of biologically independent samples (n=5). Significance was calculated using two-way ANOVA with Šidäk post-hoc test. **(G)** MDA-MB-231, MDA-MB-468 and A357 cells were cultured in cystine-free medium and treated with 500 µM GSH plus 1 µM GGsTOP and/or 2 µM liproxstatin 1 (Lip-1) for 48 hours. Relative cell number was determined by crystal violet staining. Data show mean ± SD of biologically independent samples (n=3). Significance was calculated using one-way ANOVA with Tukey post-hoc test.

**Figure S6:**
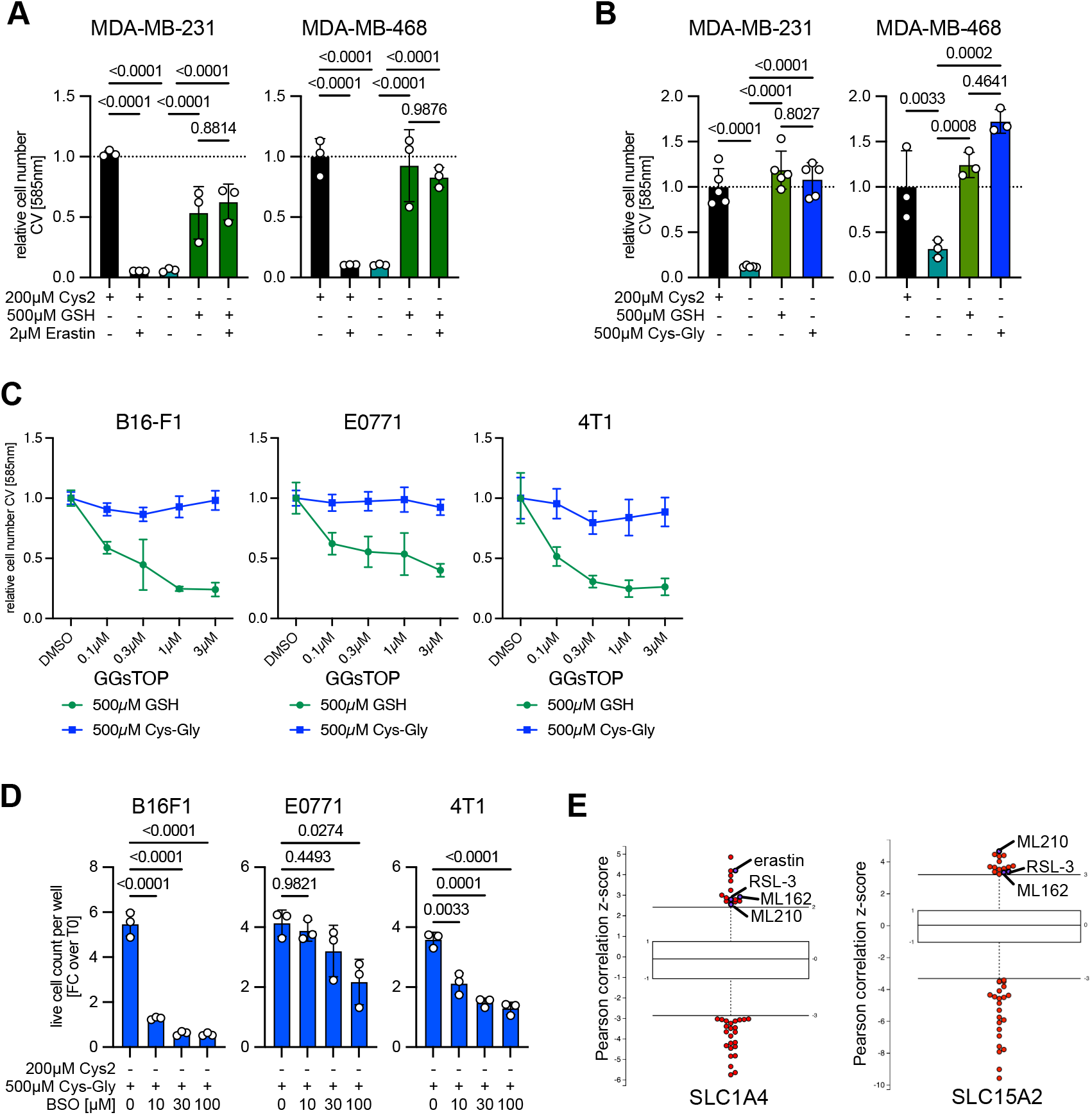
Extracellular GSH protects cancer cells against ferroptosis independent of system Xc^-^ in human cells whilst rescue by Cys-Gly might follow different uptake mechanism. **(A)** MDA-MB-231 and MDA-MB-468 cells were cultured in cystine-containing or cystine-free conditions supplemented with 500 µM GSH with or without 2 µM erastin for 96 hours. Relative cell number was determined by crystal violet staining normalized to respective controls. Data show mean ± SD of biologically independent samples (n=3). Significance was calculated using one-way ANOVA with Tukey post-hoc test on log transformed data. **(B)** The same cells were cultured in cystine-free medium with or without 500µM GSH or equimolar concentration of cysteinyl-glycine (Cys-Gly) or glycinyl-glycine (Gly-Gly) for 48 hours. Relative cell number was determined by crystal violet staining normalized to respective controls. Data show mean ± SD of biologically independent samples (MDA-MB-231 n=5; MDA-MB-468 n=3). Significance was calculated using one-way ANOVA with Tukey post-hoc test on log transformed data. **(C)** B16-F1, E0771 and 4T1 cells were cultured in cystine-free medium in the presence or absence of 500 µM GSH or 500 µM cysteinyl-glycine (Cys-Gly) and treated with the indicated concentrations of GGsTOP for 24 hours. Relative cell number was determined by crystal violet staining. Data show mean ± SD of biologically independent samples (n=6). **(D)** B16-F1, E0771 and 4T1 cells were cultured in cystine-free medium in the presence of 500 µM cysteinyl-glycine (Cys-Gly) with increasing concentrations of BSO for 48 hours. Number of live cells per well was determined by automated imaging and FC over cell count as T0 was analyzed. Data show mean ± SD of biologically independent samples (n=3). Significance was calculated using one-way ANOVA with Dunnett post-hoc test. **(E)** Correlation of drug sensitivity with the amino acid transporter SLC1A4 and the dipeptide transporter SLC15A2 expression across 860 cancer cell lines from the Cancer Therapeutics Response Portal (V2).

**Figure S7:**
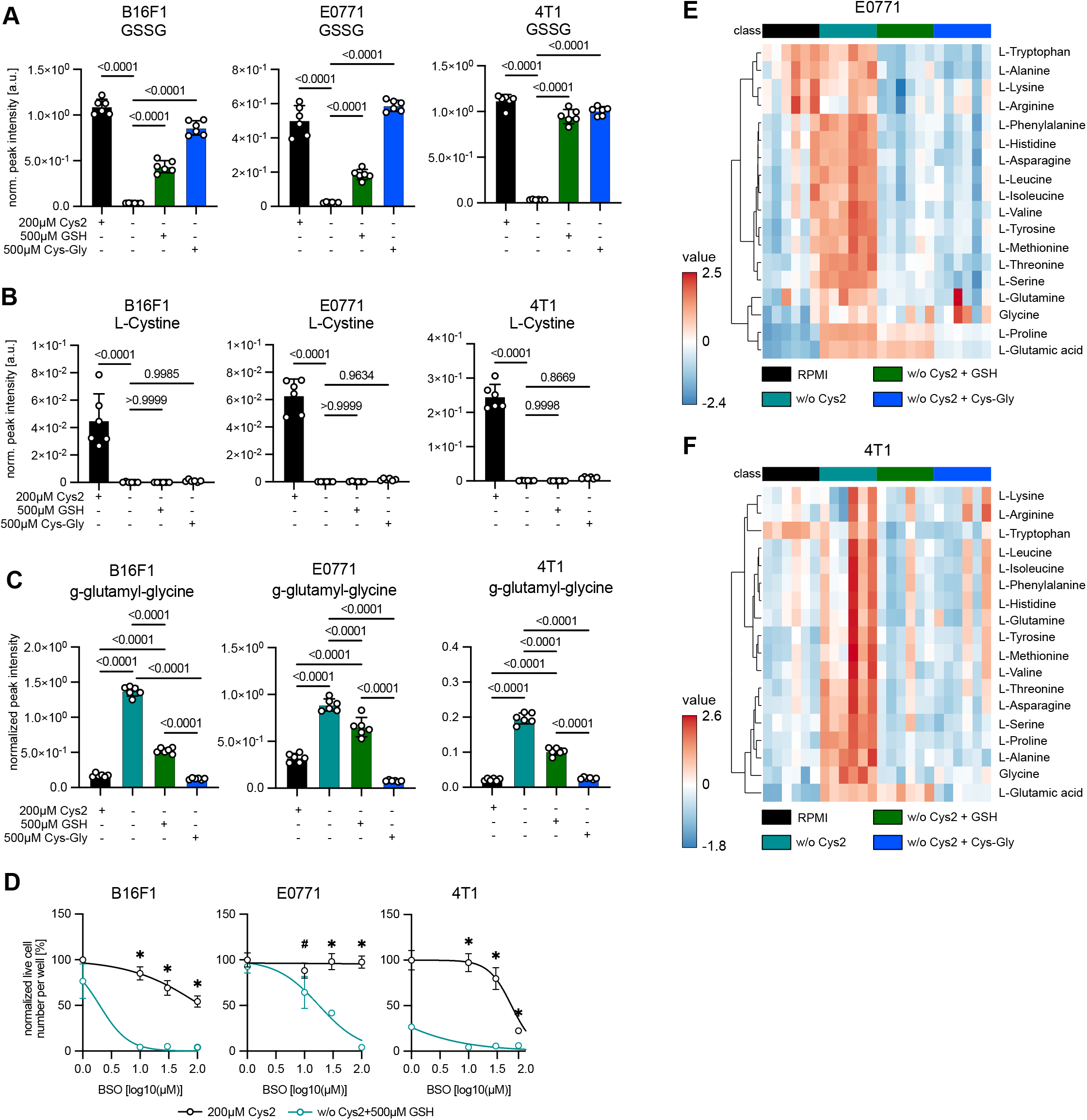
Extracellular GSH catabolism produces sufficient cysteine to restore proliferation and to circumvents amino acid accumulation in cystine-depleted conditions. **(A)** B16-F1, E0771 and 4T1 cells were cultured in RPMI (200 µM Cys2) or in cystine-free medium in the presence or absence of 500 µM GSH or 500 µM cysteinyl-glycine (Cys-Gly) for 6 hours. Graph shows levels of GSSG in cell extracts analyzed by LC-MS thiol metabolomics as this are the same samples shown in Fig 3E. Data show mean ± SD of biologically independent samples (n=6). Significance was calculated using one-way ANOVA with Tukey post-hoc test. **(B)** Intracellular levels of L-Cystine in the same cells as in shown in fig S7A. Data show mean ± SD of biologically independent samples (n=6). Significance was calculated using one-way ANOVA with Tukey post-hoc test. **(C)** Intracellular levels of g-glutamyl-glycine in the same cells as in shown in fig S7A. Data show mean ± SD of biologically independent samples (n=6). Significance was calculated using one-way ANOVA with Tukey post-hoc test. **(D)** B16-F1, E0771 and 4T1 cells were cultured in cystine-free conditions in the presence of 500 µM GSH with or without addition of BSO for 48 hours. The number of live cells per well was determined by automated counting and normalized to T0 and to respective control condition. Data show mean ± SD of biologically independent samples (n=4). Significance was calculated using one-way ANOVA with Tukey post-hoc test on log transformed data, * = *p-*value <0.0001. **(E)** Heatmap showing relative abundance of amino acids in E0771 cells treated as in fig. S7A. **(F)** Heatmap showing relative abundance of amino acids in 4T1 cells treated as in fig. S7A.

**Figure S8:**
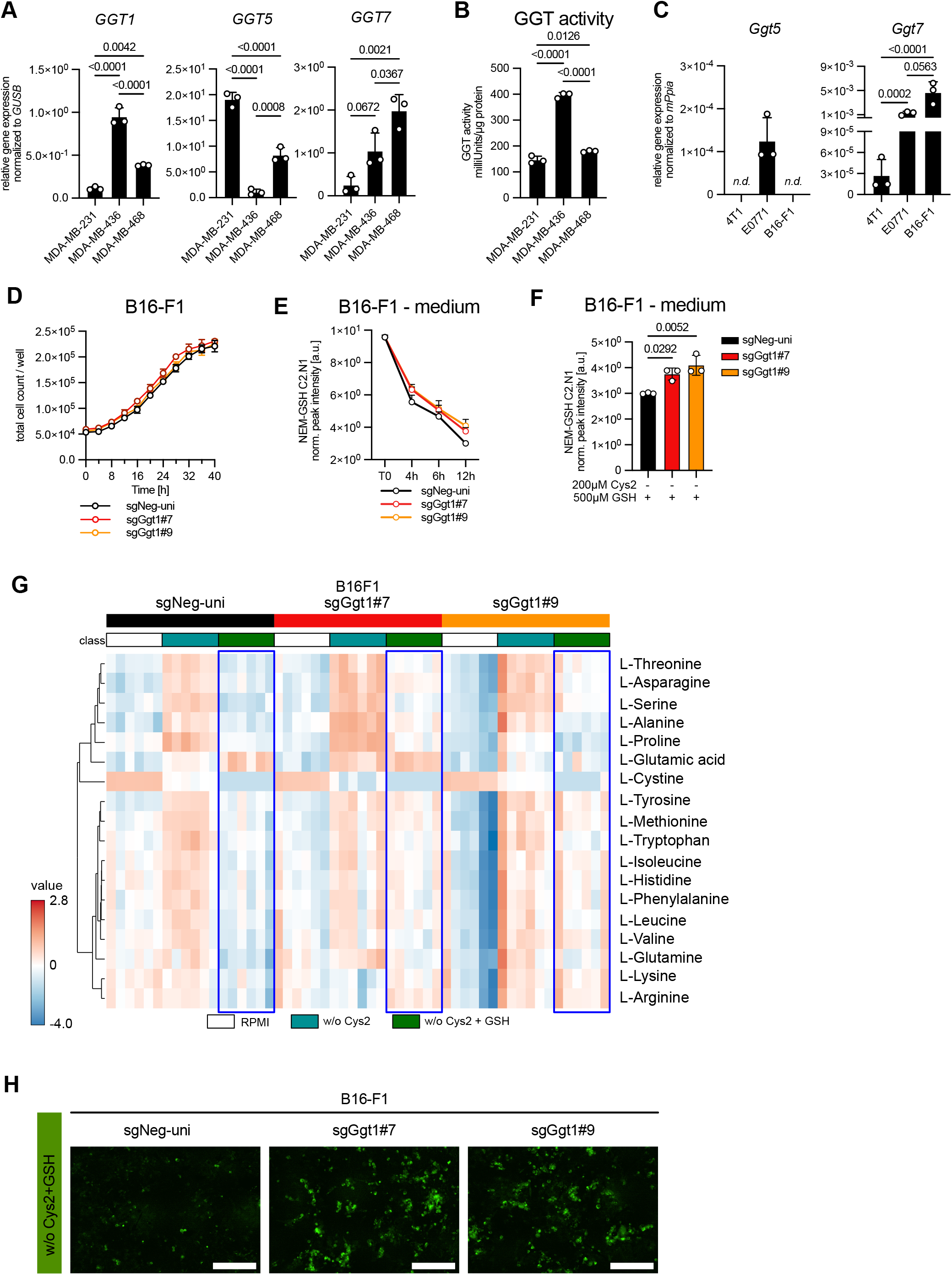
Depletion of Ggt1 attenuates extracellular GSH catabolism and rescue of amino acid accumulation in cystine-free environments. **(A)** Relative expression of *GGT1*, *GGT5* and *GGT7* mRNA in MDA-MB-231, MDA-MB-436 and MDA-MB-468 cells normalized to glucuronidase beta (*GUSB*). Data show mean ± SD of biologically independent samples (n=3). Significance was calculated using one-way ANOVA with Tukey post-hoc test. **(B)** GGT activity in the same cells determined by enzymatic assay. Data show mean ± SD of biologically independent samples (n=3). Significance was calculated using one-way ANOVA with Tukey post-hoc test. **(C)** Relative expression of *Ggt5* and *Ggt7* mRNA in 4T1, E0771 and B16-F1 cells normalised to peptidylprolyl isomerase A (*Ppia*). Data show mean ± SD of biologically independent samples (n=3). Significance was calculated using one-way ANOVA with Tukey post-hoc test on log transformed data. **(D)** Proliferation of B16-F1 control and Ggt1 knockout cells in RPMI medium. Cell number was determined by live-cell imaging. Data show mean ± SD of biologically independent samples (n=4). **(E)** B16-F1 control and Ggt1 knockout cells were cultured in RPMI or cystine-free medium in the presence of 500 µM stable isotope labelled of GSH-^13^C_2_^15^N_1_. Samples were taken after indicated time points. Relative abundance of NEM-GSH C2.N1 in medium was quantified by LC-MS thiol metabolomics. Data show mean ± SD of biologically independent samples (n=4). Significance was calculated using two-way ANOVA with Dunnett post-hoc test. **(F)** Medium levels of NEM-GSH C2.N1 at the 12h time point in the same cells as in fig. S8E. **(G)** B16-F1 control and Ggt1 knockout cells were cultured in RPMI or cystine-free medium in the presence of 500 µM GSH for 6 hours Heatmap shows relative abundance of amino acids. **(H)** B16-F1 control and Ggt1 knockout cells were cultured in cystine-free medium in the presence of 500 µM GSH for 48 hours. Representative images showing Lipid peroxidation stained by BODIPY^581/591^-C11 in cells. Quantified Data are shown in Fig 4F.

**Figure S9:**
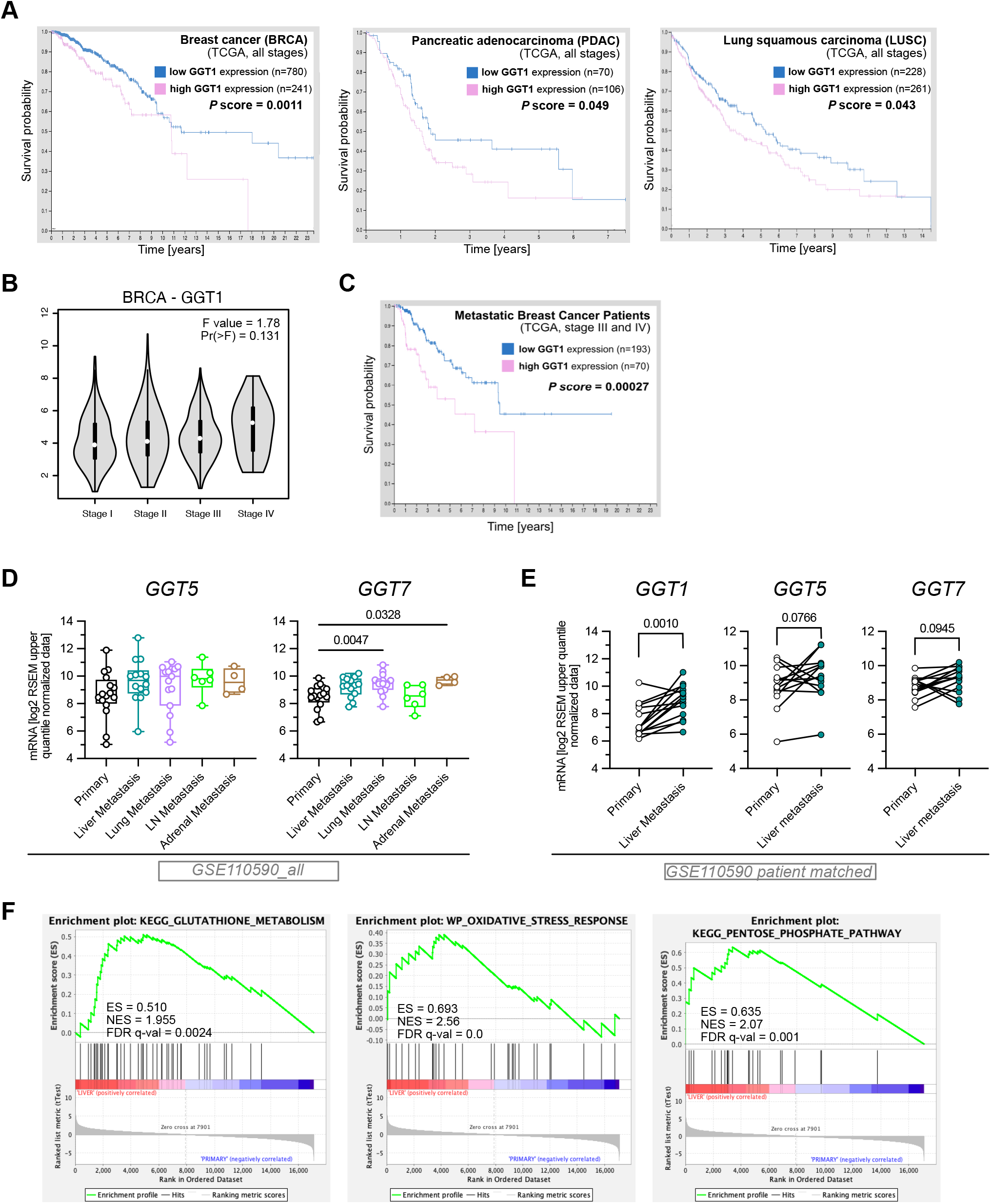
GGT1 is overexpressed in liver metastases of TNBC and relevant for patient survival. **(A)** Kaplan-Meyer curves showing association of high *GGT1* expression with poor survival in in breast (BRCA), pancreatic (PDAC) and lung squamous (LUSC) cancer derived from The Human Protein Atlas (https://www.proteinatlas.org/). **(B)** Expression of GGT1 across different stages of breast cancer analyzed using GEPIA 2 tool (http://gepia2.cancer-pku.cn/#index). **(C)** Kaplan-Meyer curves showing association of high *GGT1* expression with poor survival in metastatic breast cancer (stage III and IV). **(D)** Expression of *GGT5* and *GGT7* mRNA in a panel of TNBC primary tumors and metastases from multiple sites from GSE110590. Data are displayed as box and whisker plots indicating median, 25^th^ and 75^th^ percentile and min-max. Significance was calculated using one-way ANOVA with Dunnett post-hoc test (primary n=16, liver metastasis n=14; lung metastasis n=15; lymph node (LN) metastasis n=6; adrenal metastasis n=4). **(E)** Paired analysis of *GGT1* expression between primary tumor and matched liver metastasis from the same dataset as in (D). Significance was calculated using paired *t*-test (n=14). **(F)** Enrichment plots for the KEGG_GLUTATHIONE_METABOLISM, WP_OXIDATIVE_STRESS_RESPONSE, and KEGG_PENTOSE_PHOSPHATE_PATHWAY gene sets identified by gene set enrichment analysis (GSEA) using the C2_KEGG_LEGACY or C2_WIKIPAWTHAYS subsets comparing primary tumor and liver metastasis from GSE110590.

**Figure S10:**
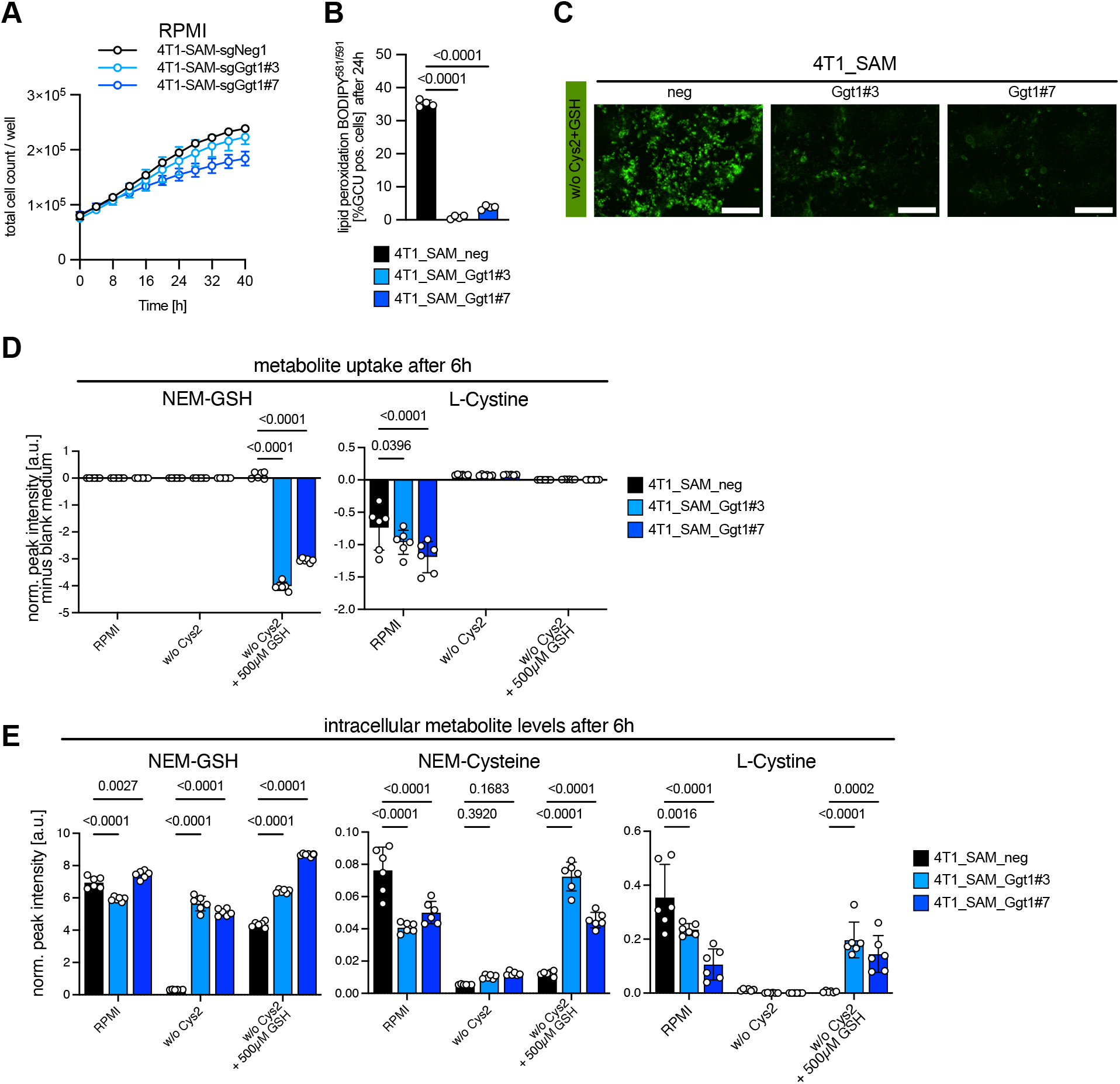
Overexpression of Ggt1 prevents leads to enhanced extracellular GSH catabolism and prevents lipid peroxidation in cystine-free environments. **(A)** Proliferation of control and Ggt1 overexpressing 4T1 cells in complete RPMI medium. Cell number was determined by live-cell imaging. Data show mean ± SD of biologically independent samples (n=4). **(B)** 4T1 control and Ggt1 overexpressing cells were cultured in cystine-free medium in the presence of 500 µM GSH for 48 hours. Lipid peroxidation was determined by BODIPY^581/591^ C11 staining. Data show mean ± SD of biologically independent samples (n=4). Significance was calculated using one-way ANOVA with Dunnett post-hoc test. **(C)** Representative images of lipid peroxidation determined by BODIPY^581/591^ C11 from fig S10B. Scale bar 200µm. **(D)** 4T1 control and Ggt1 overexpressing cells were cultured in complete or cystine-free medium in the presence or absence of 500 µM GSH for 6 hours. Uptake of GSH or L-Cystine from medium was determined by LC-MS thiol metabolomics (for detection of NEM-GSH) and calculated by subtracting respective values obtained from blank medium. Data show mean ± SD of biologically independent samples (n=6). Significance was calculated using two-way ANOVA with Tukey post-hoc test. **(E)** Intracellular levels of L-Cystine, L-Cysteine (NEM-Cysteine) and GSH (NEM-GSH) in cells cultured as in fig. S10D. Data show mean ± SD of biologically independent samples (n=6). Significance was calculated using two-way ANOVA with Tukey post-hoc test.

**Figure S11:**
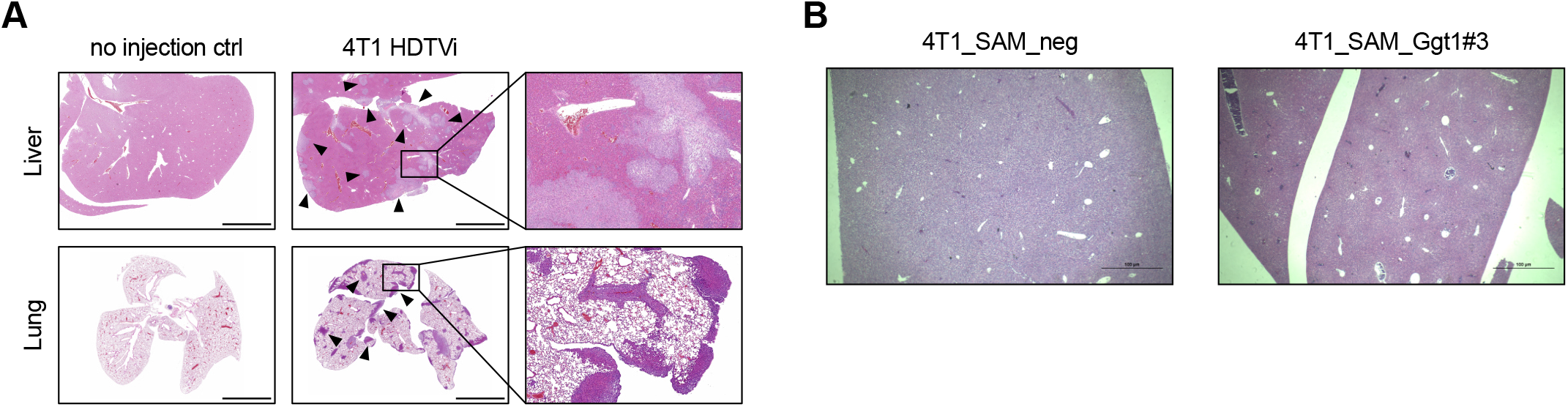
4T1 cells colonization and outgrowth in the liver and lung microenvironment after HDTVi. **(A)** Representative histological sections of matched liver and lung tissues of mice after HDTV injection of 4T1 cells (5 x 10^5^ cells per injection). Arrowheads indicate metastatic nodules visible in liver and lung sections. Scale bar 400 µm. **(B)** Representative images of liver sections from mice after orthotopic implantation and metastatic outgrowth of 4T1 control (SAM_neg) and Ggt1 overexpressing cells (SAM_Ggt1#3). No metastatic nodules were detected in the livers of mice in all groups. Livers were fixed and stained H&E. Scale bars 100 µM.

**Table S1. (separate file)**

Patient information for the tissue samples used for quantitative proteome analysis. Breast, liver and lung tissues were derived from breast cancer patients enrolled in the CATCH and COGNITION registry trials (NCT05652569 and NCT05906407).

**Table S2.**
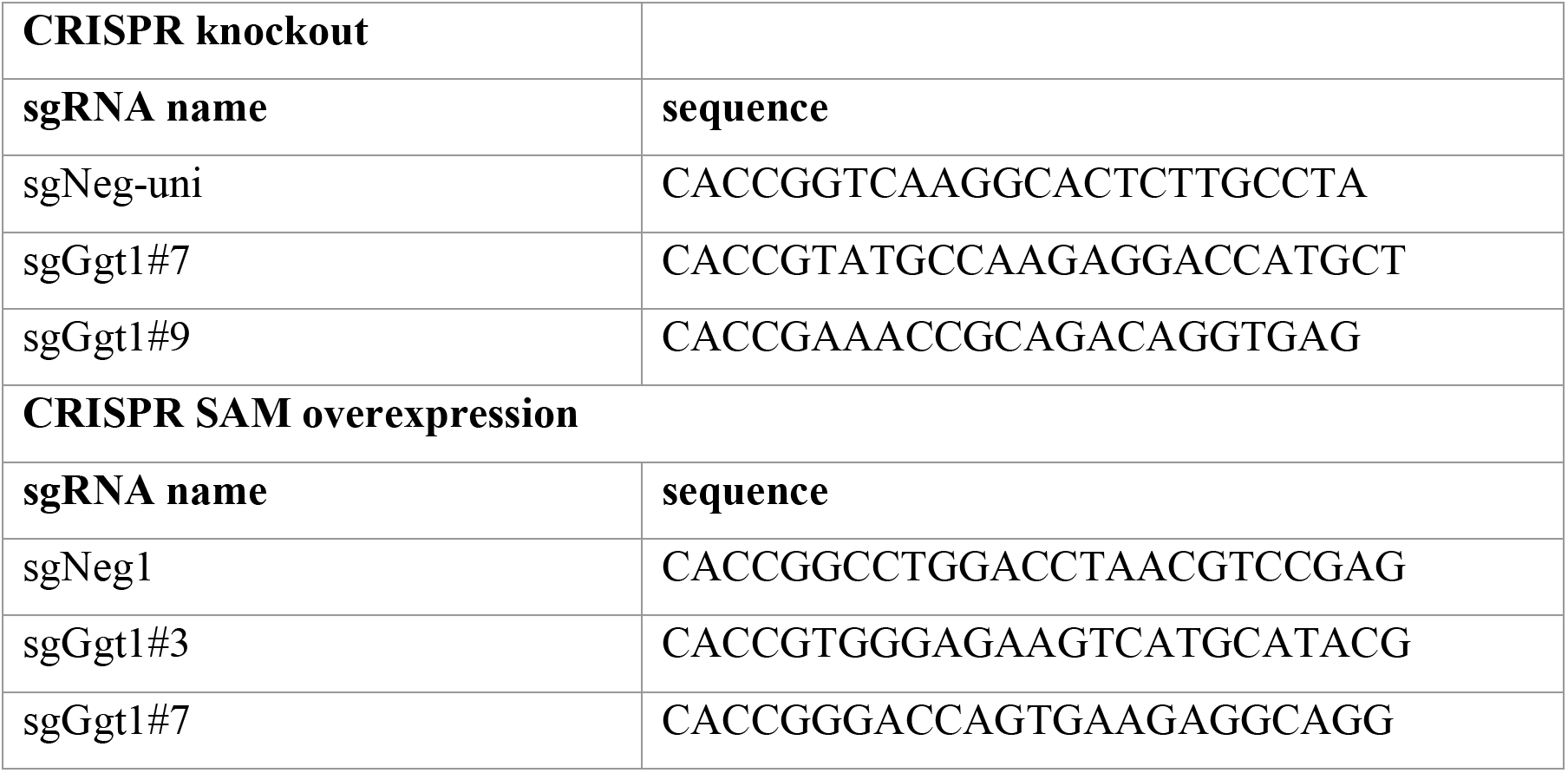
sgRNA sequences used for CRISPR gene modification.

**Table S3.**
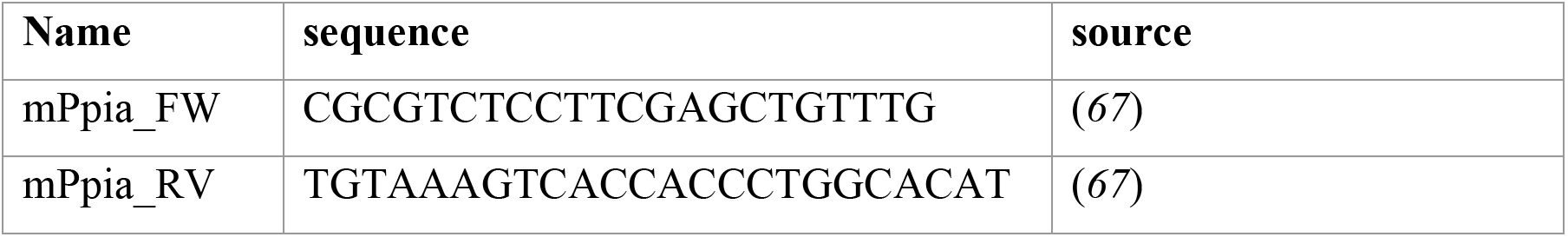
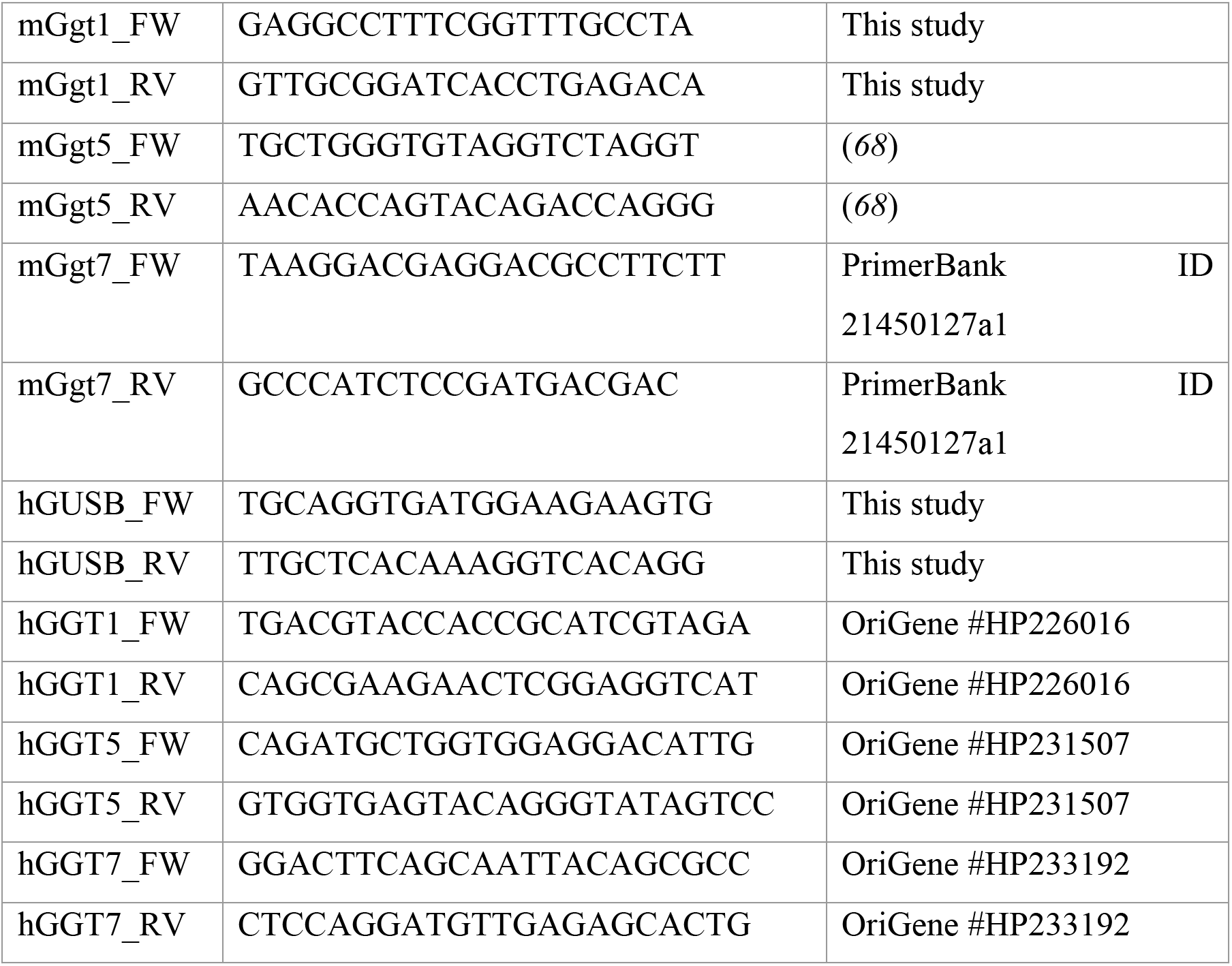
RT-qPCR primers.

**Data S1. (separate file)**

Absolute metabolite concentrations in Balb/c organ interstitial fluids and plasma determined by LC-MS using the standard addition method.

## References

1. A. W. Lambert, D. R. Pattabiraman, R. A. Weinberg, Emerging Biological Principles of Metastasis. Cell 168, 670–691 (2017).

2. G. Bergers, S. M. Fendt, The metabolism of cancer cells during metastasis. Nat Rev Cancer 21, 162–180 (2021).

3. D. X. Nguyen, P. D. Bos, J. Massague, Metastasis: from dissemination to organ-specific colonization. Nat Rev Cancer 9, 274–284 (2009).

4. A. C. Obenauf, J. Massague, Surviving at a Distance: Organ-Specific Metastasis. Trends Cancer 1, 76–91 (2015).

5. J. Massague, A. C. Obenauf, Metastatic colonization by circulating tumour cells. Nature 529, 298–306 (2016).

6. H. Basnet et al., Flura-seq identifies organ-specific metabolic adaptations during early metastatic colonization. Elife 8, (2019).

7. T. Mashimo et al., Acetate is a bioenergetic substrate for human glioblastoma and brain metastases. Cell 159, 1603–1614 (2014).

8. I. Elia et al., Breast cancer cells rely on environmental pyruvate to shape the metastatic niche. Nature 568, 117–121 (2019).

9. P. Altea-Manzano et al., A palmitate-rich metastatic niche enables metastasis growth via p65 acetylation resulting in pro-metastatic NF-kappaB signaling. Nat Cancer 4, 344–364 (2023).

10. F. Dupuy et al., PDK1-Dependent Metabolic Reprogramming Dictates Metastatic Potential in Breast Cancer. Cell Metab, (2015).

11. A. Tasdogan, J. M. Ubellacker, S. J. Morrison, Redox Regulation in Cancer Cells during Metastasis. Cancer Discov 11, 2682–2692 (2021).

12. Z. T. Schafer et al., Antioxidant and oncogene rescue of metabolic defects caused by loss of matrix attachment. Nature 461, 109–113 (2009).

13. L. Jiang et al., Reductive carboxylation supports redox homeostasis during anchorage-independent growth. Nature 532, 255–258 (2016).

14. A. Tasdogan et al., Metabolic heterogeneity confers differences in melanoma metastatic potential. Nature 577, 115–120 (2020).

15. S. J. Dixon et al., Ferroptosis: an iron-dependent form of nonapoptotic cell death. Cell 149, 1060–1072 (2012).

16. W. S. Yang, B. R. Stockwell, Synthetic lethal screening identifies compounds activating iron-dependent, nonapoptotic cell death in oncogenic-RAS-harboring cancer cells. Chem Biol 15, 234–245 (2008).

17. M. A. Badgley et al., Cysteine depletion induces pancreatic tumor ferroptosis in mice. Science 368, 85–89 (2020).

18. X. Jiang, B. R. Stockwell, M. Conrad, Ferroptosis: mechanisms, biology and role in disease. Nat Rev Mol Cell Biol 22, 266–282 (2021).

19. C. Berndt et al., Ferroptosis in Health and Disease. Redox Biology, 103211 (2024).

20. V. D. B. Bonifacio, S. A. Pereira, J. Serpa, J. B. Vicente, Cysteine metabolic circuitries: druggable targets in cancer. Br J Cancer 124, 862–879 (2021).

21. J.-Y. Min, K.-S. Chun, D.-H. Kim, The versatile utility of cysteine as a target for cancer treatment. Frontiers in Oncology Volume 12 - 2022, (2023).

22. H. F. Zhang, R. I. Klein Geltink, S. J. Parker, P. H. Sorensen, Transsulfuration, minor player or crucial for cysteine homeostasis in cancer. Trends Cell Biol 32, 800–814 (2022).

23. S. Bannai, Exchange of cystine and glutamate across plasma membrane of human fibroblasts. J Biol Chem 261, 2256–2263 (1986).

24. J. L. Parker et al., Molecular basis for redox control by the human cystine/glutamate antiporter system xc(). Nat Commun 12, 7147 (2021).

25. N. Heisterkamp, J. Groffen, D. Warburton, T. P. Sneddon, The human gamma-glutamyltransferase gene family. Hum Genet 123, 321–332 (2008).

26. M. H. Hanigan, Gamma-glutamyl transpeptidase: redox regulation and drug resistance. Adv Cancer Res 122, 103–141 (2014).

27. K. Hayashima, H. Katoh, Expression of gamma-glutamyltransferase 1 in glioblastoma cells confers resistance to cystine deprivation-induced ferroptosis. J Biol Chem 298, 101703 (2022).

28. M. H. Hanigan, B. C. Gallagher, D. M. Townsend, V. Gabarra, Gamma-glutamyl transpeptidase accelerates tumor growth and increases the resistance of tumors to cisplatin in vivo. Carcinogenesis 20, 553–559 (1999).

29. F. Hecht et al., Catabolism of extracellular glutathione supplies cysteine to support tumours. Nature 653, 933–941 (2026).

30. H. Wiig, K. Aukland, O. Tenstad, Isolation of interstitial fluid from rat mammary tumors by a centrifugation method. Am J Physiol Heart Circ Physiol 284, H416–424 (2003).

31. M. R. Sullivan et al., Quantification of microenvironmental metabolites in murine cancers reveals determinants of tumor nutrient availability. Elife 8, (2019).

32. J. R. Cantor et al., Physiologic Medium Rewires Cellular Metabolism and Reveals Uric Acid as an Endogenous Inhibitor of UMP Synthase. Cell 169, 258–272 e217 (2017).

33. M. T. Snaebjornsson et al., Targeting aldolase A in hepatocellular carcinoma leads to imbalanced glycolysis and energy stress due to uncontrolled FBP accumulation. Nat Metab 7, 348–366 (2025).

34. J. A. Combs, G. M. DeNicola, The Non-Essential Amino Acid Cysteine Becomes Essential for Tumor Proliferation and Survival. Cancers (Basel) 11, (2019).

35. L. Han, J. Hiratake, A. Kamiyama, K. Sakata, Design, synthesis, and evaluation of gamma-phosphono diester analogues of glutamate as highly potent inhibitors and active site probes of gamma-glutamyl transpeptidase. Biochemistry 46, 1432–1447 (2007).

36. K. Ortmayr, M. Schwaiger, S. Hann, G. Koellensperger, An integrated metabolomics workflow for the quantification of sulfur pathway intermediates employing thiol protection with N-ethyl maleimide and hydrophilic interaction liquid chromatography tandem mass spectrometry. Analyst 140, 7687–7695 (2015).

37. Y. P. Kang, G. M. DeNicola, Quantification and Tracing of Stable Isotope into Cysteine and Glutathione. Methods Mol Biol 2675, 51–63 (2023).

38. S. Dolma, S. L. Lessnick, W. C. Hahn, B. R. Stockwell, Identification of genotype-selective antitumor agents using synthetic lethal chemical screening in engineered human tumor cells. Cancer Cell 3, 285–296 (2003).

39. A. Basu et al., An interactive resource to identify cancer genetic and lineage dependencies targeted by small molecules. Cell 154, 1151–1161 (2013).

40. R. Dringen, B. Hamprecht, S. Broer, The peptide transporter PepT2 mediates the uptake of the glutathione precursor CysGly in astroglia-rich primary cultures. J Neurochem 71, 388–393 (1998).

41. Y. P. Kang et al., Non-canonical Glutamate-Cysteine Ligase Activity Protects against Ferroptosis. Cell Metab 33, 174–189 e177 (2021).

42. S. Wickham, M. B. West, P. F. Cook, M. H. Hanigan, Gamma-glutamyl compounds: substrate specificity of gamma-glutamyl transpeptidase enzymes. Anal Biochem 414, 208–214 (2011).

43. T. T. Bui et al., gamma-Glutamyl transferase 7 is a novel regulator of glioblastoma growth. BMC Cancer 15, 225 (2015).

44. M. Biondini et al., Metastatic breast cancer cells are metabolically reprogrammed to maintain redox homeostasis during metastasis. Redox Biol 75, 103276 (2024).

45. M. B. Siegel et al., Integrated RNA and DNA sequencing reveals early drivers of metastatic breast cancer. J Clin Invest 128, 1371–1383 (2018).

46. S. Konermann et al., Genome-scale transcriptional activation by an engineered CRISPR-Cas9 complex. Nature 517, 583–588 (2015).

47. A. Bansal et al., Gamma-Glutamyltransferase 1 Promotes Clear Cell Renal Cell Carcinoma Initiation and Progression. Mol Cancer Res 17, 1881–1892 (2019).

48. J. Li, Q. Yao, D. Liu, Hydrodynamic cell delivery for simultaneous establishment of tumor growth in mouse lung, liver and kidney. Cancer Biol Ther 12, 737–741 (2011).

49. G. Mattavelli et al., A TLR4-dependent fibroblast-monocyte axis in tumor-draining lymph nodes contributes to metastasis in triple-negative breast cancer. Immunity 58, 2830–2846 e2815 (2025).

50. G. Doglioni et al., Aspartate signalling drives lung metastasis via alternative translation. Nature 638, 244–250 (2025).

51. K. L. Abbott et al., Nutrient requirements of organ-specific metastasis in breast cancer. Nature 649, 1292–1301 (2026).

52. H. Zhang et al., 4-Hydroxynonenal induces rat gamma-glutamyl transpeptidase through mitogen-activated protein kinase-mediated electrophile response element/nuclear factor erythroid 2-related factor 2 signaling. Am J Respir Cell Mol Biol 34, 174–181 (2006).

53. S. Pandur, S. Pankiv, M. Johannessen, U. Moens, N. E. Huseby, Gamma-glutamyltransferase is upregulated after oxidative stress through the Ras signal transduction pathway in rat colon carcinoma cells. Free Radic Res 41, 1376–1384 (2007).

54. S. Reuter et al., Tumor necrosis factor alpha induces gamma-glutamyltransferase expression via nuclear factor-kappaB in cooperation with Sp1. Biochem Pharmacol 77, 397–411 (2009).

55. E. Obrador et al., gamma-Glutamyl transpeptidase overexpression increases metastatic growth of B16 melanoma cells in the mouse liver. Hepatology 35, 74–81 (2002).

56. M. Conrad et al., Cell death in cancer. Cell 189, 2322–2356 (2026).

57. J. Joung et al., Genome-scale CRISPR-Cas9 knockout and transcriptional activation screening. Nat Protoc 12, 828–863 (2017).

58. N. E. Sanjana, O. Shalem, F. Zhang, Improved vectors and genome-wide libraries for CRISPR screening. Nat Methods 11, 783–784 (2014).

59. M. R. Sullivan, C. A. Lewis, A. Muir, Isolation and Quantification of Metabolite Levels in Murine Tumor Interstitial Fluid by LC/MS. Bio Protoc 9, e3427 (2019).

60. E. Melamud, L. Vastag, J. D. Rabinowitz, Metabolomic analysis and visualization engine for LC-MS data. Anal Chem 82, 9818–9826 (2010).

61. P. Heinrich et al., Correcting for natural isotope abundance and tracer impurity in MS-, MS/MS- and high-resolution-multiple-tracer-data from stable isotope labeling experiments with IsoCorrectoR. Sci Rep 8, 17910 (2018).

62. Z. Tang, B. Kang, C. Li, T. Chen, Z. Zhang, GEPIA2: an enhanced web server for large-scale expression profiling and interactive analysis. Nucleic Acids Res 47, W556–w560 (2019).

63. S. Hofer et al., Gemcitabine and ATR inhibitors synergize to kill PDAC cells by blocking DNA damage response. Mol Syst Biol 21, 231–253 (2025).

64. F. Hamood, F. P. Bayer, M. Wilhelm, B. Kuster, M. The, SIMSI-Transfer: Software-Assisted Reduction of Missing Values in Phosphoproteomic and Proteomic Isobaric Labeling Data Using Tandem Mass Spectrum Clustering. Mol Cell Proteomics 21, 100238 (2022).

65. M. The, P. Samaras, B. Kuster, M. Wilhelm, Reanalysis of ProteomicsDB Using an Accurate, Sensitive, and Scalable False Discovery Rate Estimation Approach for Protein Groups. Mol Cell Proteomics 21, 100437 (2022).

66. C. Tacconi et al., CD169(+) lymph node macrophages have protective functions in mouse breast cancer metastasis. Cell Rep 35, 108993 (2021).

67. K. J. Veazey, M. C. Golding, Selection of stable reference genes for quantitative rt-PCR comparisons of mouse embryonic and extra-embryonic stem cells. PLoS One 6, e27592 (2011).

68. Y. Zhang et al., Gamma-glutamyl transferase 5 overexpression in cerebrovascular endothelial cells improves brain pathology, cognition, and behavior in APP/PS1 mice. Neural Regen Res 20, 533–547 (2025).

